# 16S ribosomal RNA modification drives transcript-specific translation efficiency

**DOI:** 10.64898/2026.04.20.719615

**Authors:** Zachory M. Park, Christina R. Savage, Amanda R. Decker-Farrell, Chin-Hsien Tai, Tapan K. Maity, Weiming Yang, Lisa M. Jenkins, Kumaran S. Ramamurthi

**Affiliations:** Laboratory of Molecular Biology, National Cancer Institute, National Institutes of Health, Bethesda, MD, USA; Laboratory of Cell Biology, National Cancer Institute, National Institutes of Health, Bethesda, MD, USA

**Keywords:** SpoIVA, SpoVM, Initiation, Elongation, stationary phase, 30S subunit

## Abstract

Bacterial ribosomal RNAs (rRNAs) are decorated with conserved nucleotide modifications, but the functionality of these modifications is often underexplored. MraW (RsmH) is a 16S rRNA methyltransferase that fine-tunes ribosomal function. We identified a loss-of-function allele in *mraW* that corrected a late-stage sporulation defect in *Bacillus subtilis* by bypassing a key sporulation checkpoint via altered translational regulation. Purified ribosomes isolated from Δ*mraW* cells displayed a ∼2-fold decrease in translation efficiency; *in vivo*, Δ*mraW* cells produced decreased levels of the sporulation checkpoint protein CmpA. This regulation was mediated by sequences from the 5’ untranslated region and the coding sequence of *cmpA*, which form a step-loop structure that occlude early codons of the mRNA. Proteomic analysis revealed that MraW directly or indirectly regulates the production of multiple proteins, some of which form similar structural elements as the *cmpA* transcript. We propose that MraW modification of 16S rRNA enhances translation efficiency in general, and that specific transcripts, whose gene products are likely required in limiting quantities, have evolved structural features that act as a regulatory mechanism to govern protein levels. This type of regulation may be most apparent in bacteria which exhibit uncoupled transcription and translation.

**HIGHLIGHTS:**

- A conserved 16S rRNA modification enhances translation of structured mRNAs
- Early mRNA stem-loops impose translational control dependent on ribosome modification
- mRNA structure and rRNA modifications likely co-evolved to fine-tune protein dosage

## INTRODUCTION

Ribosomes are composed of myriad proteins and three RNA species called ribosomal RNAs (rRNA) ^1–5^ that synthesize proteins from mRNA templates. In bacteria, the active ribosome is composed of the 30S small subunit, 50S subunit, and various translation factors that participate in initiation, elongation, and eventual termination of translation ^1,3–5^. The essentiality of translation has made it a successful target for antibiotics, but many aspects regarding target specificity during translation remain underexplored. For example, instead of cells harboring uniform populations of ribosomes, studies have suggested that ribosomes in a cell may be differentially modified, and that this ribosomal heterogeneity may be instrumental in the ability of bacteria to rapidly respond to changing environments ^6–12^. In any case, chemical modifications of the ribosome may represent a mechanism that dictates protein synthesis levels of key factors during stress responses and developmental programs.

One common ribosomal modification target is rRNAs. The 16S rRNA (in the 30S subunit) and 23S rRNA (in the 50S subunit) contain widely conserved nucleotide modifications that impact ribosome assembly, translation fidelity, and antimicrobial resistance ^13–17^. Removal of any single modification typically does not impact cell survival, suggesting that these modifications have more nuanced or condition-specific roles in modulating ribosome activity ^13^. One such rRNA modification is the dual methylation of C1402 in the 16S rRNA. This modification is present in almost all bacteria and consists of methylation at the N4 and 2’O positions (m^4^Cm), catalyzed by the methyltransferases MraW (RsmH) and RsmI, respectively ^5,13,18–21^. Although deletion of either *mraW* or *rsmI* results only in minor growth defects in *Escherichia coli* ^20,22^ and *Bacillus subtilis* ^23^ in laboratory media, it does result in attenuated virulence in *E. coli* O157:H7 ^24^ and *Staphylococcus aureus* ^25^. Additional work in *E. coli* showed that loss of MraW or RsmI impaired start codon selection by increasing mis-initiation at “AUU” start codons ^20^. Consistent with this in vivo observation, structural studies revealed that the methylation at N4 of C1402 engages in direct interactions with the phosphate backbone of the mRNA start codon and participates in interactions that likely stabilize the P-site of the ribosome ^5,20^. Despite the conservation of C1402 and widespread conservation of the methyltransferases responsible for its modification, the absence of a strong growth phenotype upon loss of these modifications suggests a condition- or transcript-specific role for C1402 methylation.

Previous studies have indicated that, in addition to the well-studied regulation of transcription ^26,27^, regulation of translation also ensures the proper completion of spore formation (sporulation) in the Gram-positive bacterium *B. subtilis* ^28^. The proper initiation of sporulation itself requires translational regulation via the activity of translation elongation factors that enhance the ability of ribosomes to synthesize proteins with polyproline stretches ^29^. While proline codons themselves are in general difficult for the ribosome to decode, polyproline tracts augment this problem and can lead to ribosome stalling. Accordingly, several factors have evolved to aid the ribosome in synthesizing proteins with these sequences ^29–32^.

Sporulation is a developmental program that transforms a vegetative cell into a dormant spore when the cell senses extreme nutrient depletion ^33,34^ (Fig. 1A); spores then germinate and continue vegetative growth when favorable environmental conditions resume ^35^. Spores are highly resistant to environmental insults ^36^, and this resistance is partially mediated by two cell surface structures that encase the spore: a thick proteinaceous “coat” and a peptidoglycan “cortex” that is sandwiched between two membranes ^37^. The basement layer of the spore coat is composed of the ATPase SpoIVA ^38–40^, a cytoskeletal protein that forms static homopolymers upon ATP hydrolysis which serves as a platform for the subsequent recruitment and assembly of all other coat proteins ^41–43^. Disrupting the polymerization of SpoIVA impairs not only coat assembly, but also cortex assembly via activation of the coat assembly checkpoint ^44^ that contains at least two independent pathways: the SpoVID-mediated coat assembly monitoring system ^45,46^ and the CmpA-mediated degradation system which promotes the proteolysis of defective SpoIVA subunits that do not polymerize ^47^. Together, both systems ensure that spores with mis-assembled coats do not progress to the cortex assembly step and can therefore be cleared from the population ^48^.

**Figure 1.**
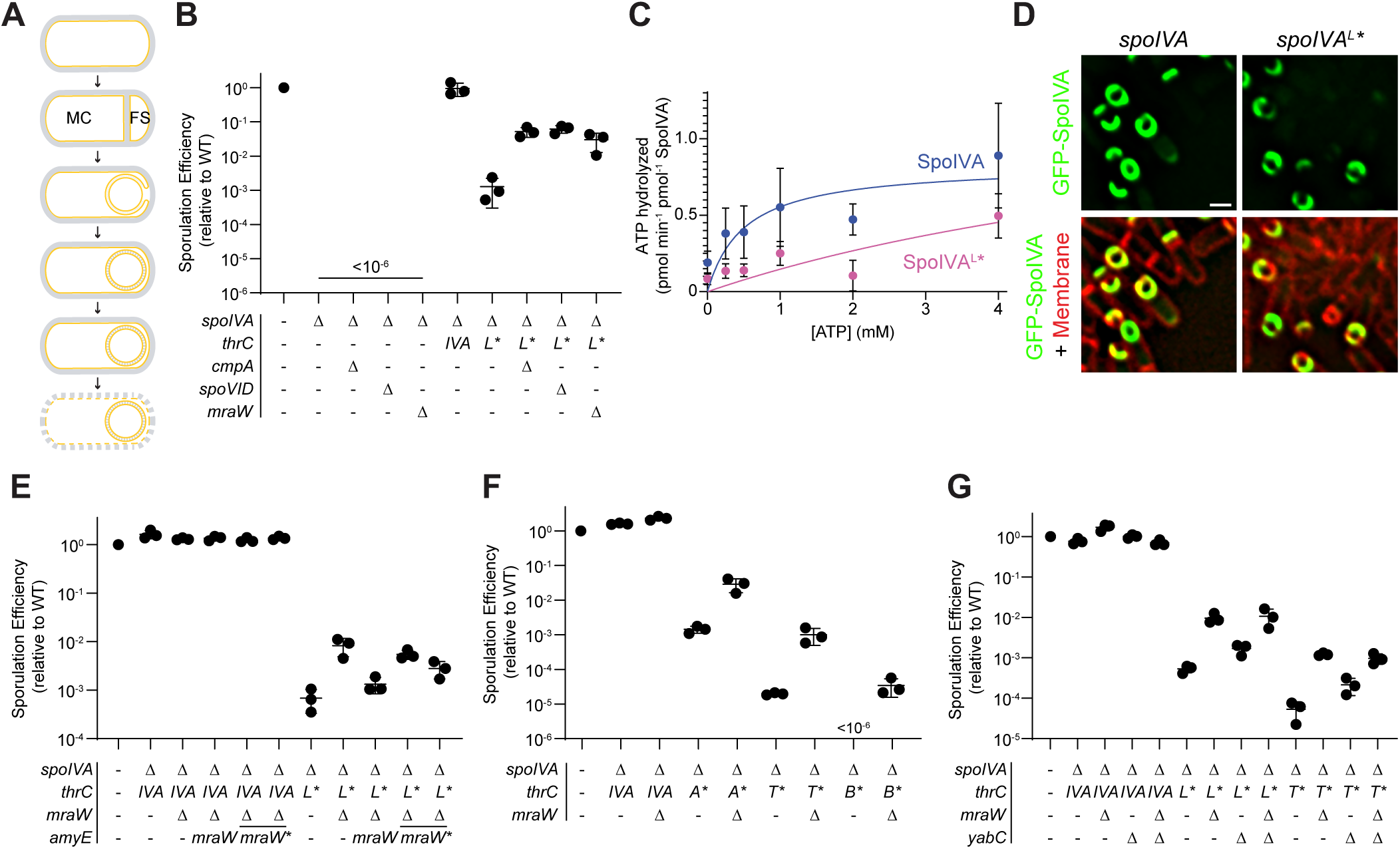
Deletion of *mraW* and its partner rRNA modifying enzyme suppress mutant SpoIVA sporulation defects. (A) Schematic of the major steps of sporulation. Membranes depicted in yellow; cell wall, gray. MC: mother cell; FS: forespore. (B) Sporulation efficiencies (determined as resistance to wet heat at 80 °C for 20 min) of strains harboring the indicated alleles of *spoIVA* in the presence or absence (Δ) of *cmpA*, *spoVID*, or *mraW*. *thrC* is a chromosomal locus into which the indicated allele of *spoIVA* is integrated to complement the *spoIVA* deletion. Strains: PY79, KP73, IT895, JPC282, ZP44, KR394, ZP47, ZP50, ZP53, and ZP56. Bars represent mean (n = 3 biological replicates); errors: S.D. (C) ATPase activity (measured as generation of free phosphate) of purified WT SpoIVA (blue) and SpoIVA^L59P^ (pink). Data were fit to the Michaelis-Menten enzyme saturation model. Data points represent mean (n = 3 independent trials); errors: S.E. (D) Subcellular localization of GFP-SpoIVA (left) and GFP-SpoIVA^L59P^ (right) using fluorescence microscopy. Top row: fluorescence from GFP (green); bottom row: overlay, GFP and membranes visualized using FM 4-64 (red). Scale bar: 1 µM. Strains KR160 and ZP35. (E-G) Sporulation efficiencies of cells harboring (E) *spoIVA*^L^* in the presence or absence of *mraW* or *mraW\** (D52A and D100A, respectively), (F) the indicated allele of *spoIVA* in the presence or absence of *mraW*, or (G) the indicated allele of *spoIVA* in the presence or absence of *yabC*. *amyE* is a chromosomal locus into which the indicated allele of *mraW* is integrated to complement the *mraW* deletion. Strains: (E) PY79, KR394, ZP77, KR367, ZP62, JPC221, ZP65, JB103, ZP59; (F) PY79, KR394, ZP77, ZP134, ZP231, ZP233, ZP47, ZP56, ZP137, ZP239, and ZP231; (G) PY79, KR394, ZP77, ZP221, ZP287, ZP47, ZP56, ZP215, ZP283, JPC221, ZP65, ZP249, and ZP285. Bars represent mean (n = 3 biological replicates); errors: S.D.

Here, we report that inactivation of *mraW* bypasses the coat assembly checkpoint and suppresses sporulation defects associated with defective alleles of *spoIVA*. We show that ribosomes isolated from Δ*mraW* cells are slightly defective for translation in vitro. In vivo, MraW modification of 16S rRNA facilitates the efficient translation of *cmpA* mRNA. We observed that *cmpA* mRNA harbors a conserved stem-loop structure that necessitates MraW modification of C1402 for efficient translation. Proteomic analysis revealed other MraW-dependent transcripts in stationary phase that also harbor a similar stem-loop structure. We propose that certain mRNA transcripts, particularly those that encode for inhibitory proteins like CmpA that are only required in limiting quantities, evolved stem-loop structures that are intrinsically difficult to translate and that require methylation of C1402 in 16S rRNA that serves as a regulatory mechanism to modulate cellular protein levels.

## RESULTS

### SpoIVA^L59P^ is defective for ATP hydrolysis and sporulation

The major structural component of the spore coat basement layer in *B. subtilis* is a structural protein called SpoIVA ^38^. SpoIVA is a TRAFAC family NTPase that hydrolyzes ATP to drive its irreversible polymerization to form a platform at the forespore surface on which subsequent spore coat proteins assemble ^42,43^. The SpoIVA^L59P^ variant was previously identified using PCR mutagenesis as a variant that confers a sporulation defect ^49^. Whereas other well-characterized loss-of-function SpoIVA variants harbored disruptions in the nucleotide binding site of SpoIVA, position L59 is located far away from the active site. We therefore sought to explore the defect associated with the L59P substitution. Cells harboring *spoIVA*^L59P^ (hereafter, simply “*spoIVA*^L^*”) displayed a ∼1000-fold reduction in sporulation efficiency compared to WT (Fig. 1B), despite the variant being produced at similar levels as the WT SpoIVA (Fig. S1A). We next purified recombinant SpoIVA^L^* and measured ATP hydrolysis at increasing nucleotide concentrations in vitro to produce saturation curves that revealed the substrate turnover rate (*k*_cat_). Compared to WT SpoIVA, which displayed a *k*_cat_ of 0.83 pmol min^-1^ pmol^-1^ SpoIVA ^40^, SpoIVA^L^* did not specifically hydrolyze ATP, as evidenced by the failure of the reaction to reach saturation and display Michaelis-Menten kinetics (Fig. 1C). Examination of the subcellular localization of SpoIVA^L^*-GFP revealed that, similar to SpoIVA variants harboring disruptions in the nucleotide binding site, SpoIVA^L^*-GFP displays a slight localization defect in that it fails to uniformly polymerize around the forespore and instead localizes as caps on the mother cell-distal and -proximal sides of the forespore (Fig. 1D) ^45^. We conclude that the L59P substitution likely causes a structural defect in SpoIVA that disrupts its ability to hydrolyze ATP and therefore inhibits sporulation by activating the coat assembly checkpoint ^44^.

### Deletion of *mraW* suppresses several *spoIVA* mutants

We took advantage of the severe sporulation defect of *spoIVA*^L^* to isolate spontaneous mutants that would correct this defect. Cells were induced to sporulate, after which nonsporulating and poorly sporulating cells were killed by exposure to 80 °C for 20 mins. Surviving cells were enriched by repeated dilution in fresh sporulation media where they could germinate and re-sporulate. Whole genome sequencing revealed a guanidyl insertion in the *mraW* (*rsmH*) gene at position 597, resulting in a frameshift that altered the next three amino acids and introduced a premature stop codon that would truncate the protein by 109 residues (approximately one third of the protein). To test if this mutation resulted in a loss of function in MraW, we examined the sporulation efficiencies of cells harboring *spoIVA*^L^* in the presence and absence of *mraW*. Deletion of *mraW* did not affect the sporulation efficiency of otherwise WT cells (Fig. 1E, lane 3) but did increase the sporulation efficiency of cells harboring *spoIVA*^L^* ∼10-fold (Fig. 1B, lane 10, Fig. 1E, lane 9). Suppression of the *spoIVA^L^\** defect was reversed upon complementation of *mraW* at an ectopic genomic locus (Fig. 1E, lane 9), indicating that the suppression is due to the loss of *mraW*. Deletion of *mraW* also partially corrected the sporulation defects of other *spoIVA* alleles that disrupted residues in the nucleotide binding pocket, including disruption of the SpoIVA Walker A motif (*spoIVA*^A^*), sensor threonine (*spoIVA*^T^*), and Walker B motif (*spoIVA*^B^*) (Fig. 1F), indicating that the suppressor was not allele-specific.

MraW is a methyltransferase that modifies 16S ribosomal RNA by methylating the conserved cytidine at position 1402 in *E. coli* that sits in the ribosomal peptidyl tRNA binding site (P-site) ^20^. To determine if loss of the methyltransferase activity specifically of MraW suppresses the sporulation defect caused by defective *spoIVA* alleles, we identified putative residues involved in binding S-adenyl-methionine guided by the co-crystal structure of *E. coli* MraW ^19^ and substituted those residues with Ala. Disruption of either D52 or D100 did not affect accumulation of the protein in vivo (Fig. S1B), but did partially suppress the sporulation defect caused by *spoIVA*^L^* (Fig. 1E, lanes 10-11; *mraW*\* represents D52A and D100A variants, respectively). Whereas MraW methylates the cytosine base of C1402 ^19,20^, C1402 in bacteria is also methylated on the deoxyribose moiety by RsmI ^20^, which in *B. subtilis* is predicted to be encoded by *yabC*. Deleting *yabC* also partially suppressed the sporulation defects of *spoIVA*^L^* and *spoIVA*^T^*, albeit not to the same extent as deleting *mraW* (Fig. 1G). These varying impacts of deleting either *mraW* or *yabC* are consistent with the epistatic effects of these mutations reported for *E. coli* (wherein the double mutant phenocopies the deletion of *rsmH*/*mraW* alone) ^20^. Together, the data suggest that the loss of methylation of the 16S rRNA permits increased sporulation despite the presence of coat assembly defects.

### MraW impacts translation in vitro

Several studies have reported the effects of deleting *mraW* on the expression of various constructs in vivo in *E. coli* ^20^, but a direct effect on translation has not been reported. To determine if the loss of MraW directly affects the functionality of the ribosome, we purified ribosomes from either WT or Δ*mraW* cells and assayed their ability to produce GFP from a transcript expressed from the T7 promoter in a combined transcription-translation reaction. WT ribosomes synthesized GFP, but ribosomes from Δ*mraW* produced > 2-fold less GFP (Fig. 2A). This reduction in activity was not due to reduced levels of ribosomes, since ribosomes purified from Δ*mraW* cells contained similar quantities of rRNA (Fig. 2B) and total protein (which displayed a similar banding pattern; Fig. 2C), suggesting that reduced GFP levels by the Δ*mraW*-derived ribosomes were due to compromised translation efficiency. The relatively large decrease in translation efficiency suggested that deleting *mraW* may result in a growth defect in vivo. However, we observed no difference in growth rate between WT and Δ*mraW* in two different rich media and one more defined medium (Fig. S2A-C). We therefore suspect that the in vitro translation conditions present a stringent environment where a strong defect can be observed, whereas rapid growth in nutrient-rich conditions largely masks these effects. Consistent with this hypothesis, we observed similar production of GFP from a construct expressed by the T7 promoter in vivo in the presence and absence of MraW (Fig. S2D). The results therefore indicate that loss of methylation of the 16S rRNA by MraW abrogates the ability of ribosomes to efficiently translate, especially under growth-limiting conditions.

**Figure 2.**
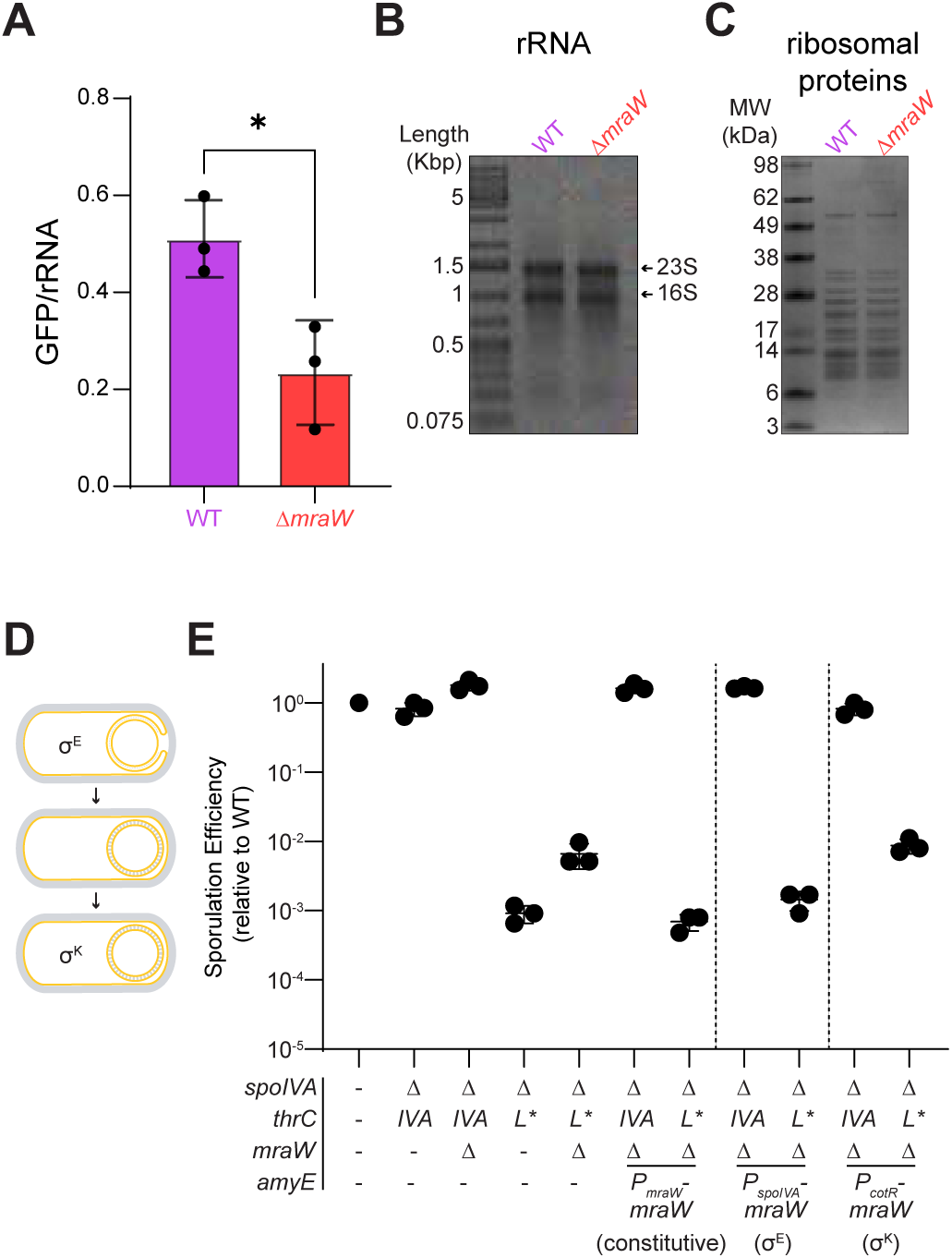
MraW-modified ribosomes promote robust translation *in vitro* and are required early during sporulation to ensure spore quality control. (A) GFP fluorescence generated from an in vitro transcription/translation assay using ribosomes purified from WT or Δ*mraW B. subtilis* (strains PY79 or ZP1) after 3 hours of growth in LB 10/10/10 media. Bars represent mean (n = 3 independent trials using 3 separate sets of ribosomes purified from each strain); errors: S.D. (B) Non-denaturing agarose gel of ribosomes purified from WT and Δ*mraW* stained with propidium iodine. Predicted bands for the 23S and 16S rRNA based on electrophoretic mobility are labeled. Lane 1: DNA size marker with length (kilobase pairs) indicated to the left. (C) Coomassie stained PAGE gel of the same ribosomal samples from (B). Lane 1: molecular weight markers with sizes (kDa) indicated to the left. (D) Schematic depicting the timing of expression of two mother cell specific sigma (σ) factors, σ^E^ and σ^K^. σ^E^ is the first mother cell specific sigma factor and σ^K^ is the final mother cell specific sigma factor. (E) Sporulation efficiencies of indicated strains that express *mraW* constitutively or under control of a σ^E^-controlled promoter (*P_spoIVA_*) or σ^K^-controlled promoter (*P_cotR_*). Strains: PY79, KR394, ZP77, ZP47, ZP56, ZP134, ZP137, ZP89, ZP83, ZP107, and ZP104.

### Early presence of MraW during sporulation is required to activate the coat assembly checkpoint

Given the relatively reduced translation efficiency of ribosomes that were unmodified by MraW, we reasoned that deletion of *mraW* may suppress the sporulation defect caused by SpoIVA variants by specifically impacting the translation of a sporulation factor (likely a negative regulator of sporulation). To perform a candidate-based approach to identify this factor, we first assessed the point during the sporulation program that MraW is required to block sporulation in the presence of a defective SpoIVA variant. SpoIVA is produced in the mother cell shortly after asymmetric division, under control of the σ^E^ sigma factor (Fig. 2D). We therefore expressed *mraW* under control of either a σ^E^-driven promoter, or a promoter driven by a late mother cell sigma factor (σ^K^; Fig. 2D) and monitored if either construct would complement the deletion of *mraW*. Expression of *mraW* from a σ^E^-driven promoter complemented the deletion of *mraW* similar to the complementation of Δ*mraW* by *mraW* under its native promoter (Fig. 2E, lanes 6-9), but expression of *mraW* from a σ^K^-driven promoter did not (Fig. 2E, lanes 10-11). We therefore conclude that MraW is required relatively early during sporulation to block cells harboring defective SpoIVA variants from forming heat-resistant spores.

### MraW is required for the efficient translation of CmpA

There are two known negative regulators of sporulation that participate in the coat assembly checkpoint that prevents cells from proceeding through the sporulation program if the cell harbors a defective SpoIVA variant: SpoVID ^45^ and CmpA ^47^. We therefore investigated, using β-galactosidase fusions, if deletion of *mraW* affects the production of either protein. Expression of *spoVID* was not appreciably affected by deletion of *mraW* (Fig. S3A-B). However, a translational fusion to the *cmpA* open reading frame (ORF) revealed reduced translation of *cmpA* over time in the absence of *mraW* in cells producing a defective allele of *spoIVA* (Fig. 3A) or in otherwise WT cells (Fig. 3C). This effect of MraW was not observed when we employed a transcriptional fusion to the *cmpA* promoter only (Fig. 3B, 3D). Moreover, the in vivo half-life of CmpA-LacZ was similar in a spectinomycin chase experiment in the presence and absence of *mraW*, as was the expression of *lacZ* alone when fused to a constitutive promoter (Fig. S3C-D), indicating that the turnover of CmpA and the synthesis of LacZ alone were unaffected by the absence of MraW. Finally, deletion of *mraW* also reduced production of CmpA-GFP from an overexpressed transcript encoding a translational fusion of *cmpA*-*gfp* driven by a constitutive promoter (instead of the relatively weak *P*_cmpA_ promoter), suggesting that MraW specifically affects the translation of *cmpA* (Fig. S3E). Overexpression of *cmpA* from a constitutive promoter reduces sporulation efficiency even in an otherwise WT cell ^47^ (Fig. 3E, lanes 2 and 6). This overexpression effect is more pronounced in the presence of a defective *spoIVA* allele (Fig. 3E, lane 10). However, consistent with the previous observation, deletion of *mraW* also suppressed this sporulation defect arising from *cmpA* overexpression, either in the presence of WT SpoIVA (Fig. 3E, lane 3, 7) or in the presence of SpoIVA^T^* (Fig. 3E, lane 11). We therefore hypothesized that deletion of *mraW*, and the resulting loss of 16S rRNA methylation, negatively impacts translation of the *cmpA* mRNA, which in turn suppresses the sporulation defect of defective SpoIVA variants.

**Figure 3.**
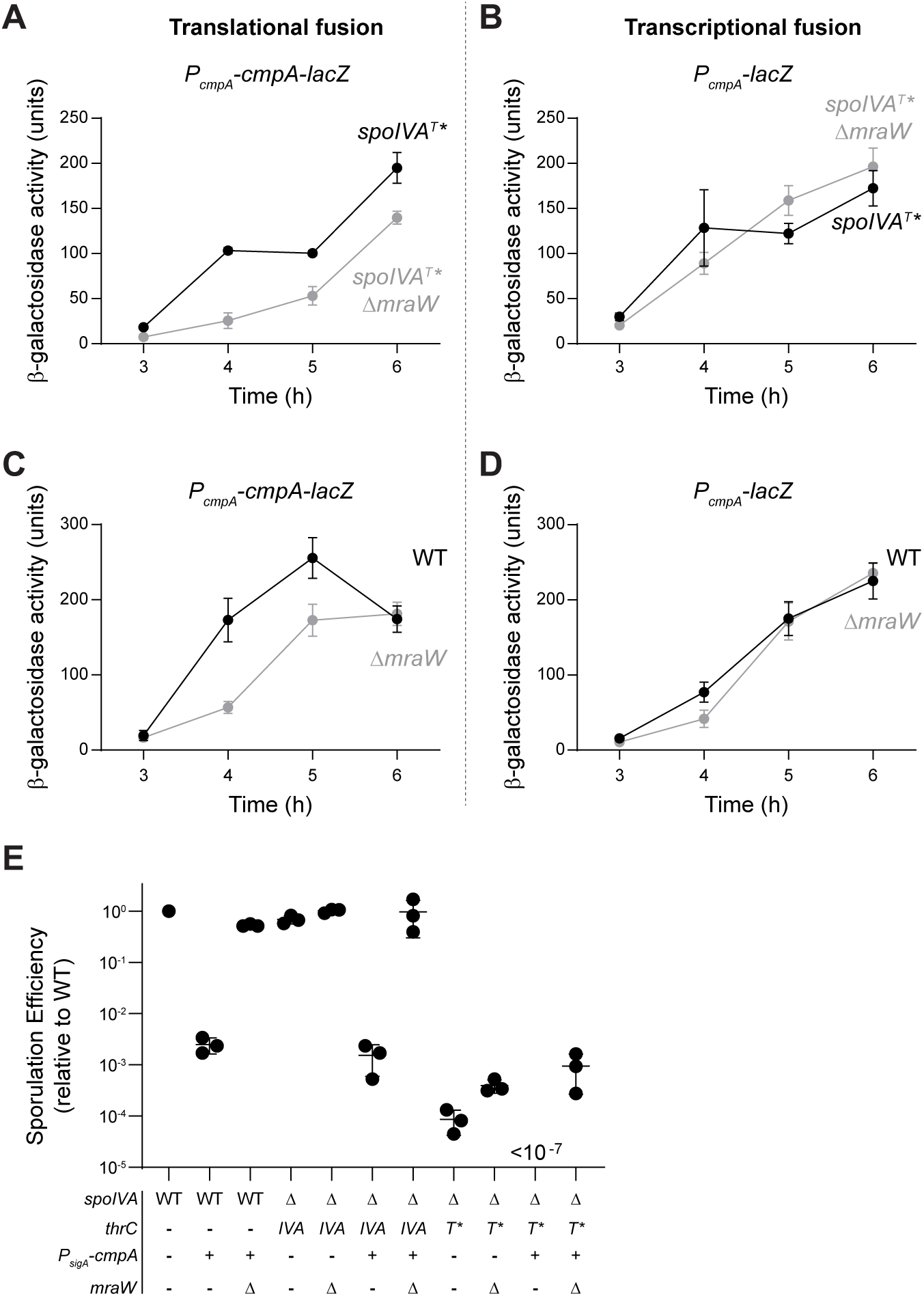
CmpA translation, but not transcription, is reduced in Δ*mraW*. (A-D) β-galactosidase activity arising from (A, C) translational fusions of the *cmpA* ORF to *lacZ* expressed from the *cmpA* promoter or (B, D) transcriptional fusions of the *cmpA* promoter fused directly to *lacZ* in cells harboring (A-B) *spoIVA*^T^* or (C-D) WT *spoIVA*, in the (black) presence or (gray) absence of *mraW* at indicated time points after induction of sporulation. Data points represent mean (n = 3 biological replicates; errors: S.D.). Strains: ZP385, ZP389, ZP383, ZP387, SE230, ZP313, SE222, and ZP311. (E) Sporulation efficiencies of strains in the presence of *cmpA* that is constitutively produced (*P_sigA_-cmpA*) harboring the indicated allele of *spoIVA* in the presence or absence of *mraW*. Bars represent mean (n = 3 biological replicates); errors: S.D. Strains: PY79, IT686, ZP116, KR394, ZP77, ZP315, ZP321, JPC221, ZP65, ZP317, and ZP319.

### MraW regulates CmpA protein levels via a feature of the cmpA mRNA sequence

Next, we investigated if an element within the *cmpA* transcript is responsible for the MraW-mediated regulation of CmpA protein levels. First, we truncated the *cmpA* ORF fused to lacZ to identify the minimal region responsible for this regulation. Sequential 3’ truncations of the 37-codon-long *cmpA* ORF showed that MraW-mediated regulation persisted when the 5’ untranslated region was fused to just the first 10 codons of *cmpA* (Fig. 4A-D). To test if specific codon usage was important for the Δ*mraW* effect, we fused the upstream untranslated region and codons 1-20 to *lacZ*, introduced a -1 frameshift immediately after the *cmpA* start codon and corrected the reading frame by adding a nucleotide at the fusion to *lacZ* to restore the proper reading frame (Fig. 4E). This disruption largely preserved the nucleotide sequence of *cmpA*^1-20^ but disrupted the amino acid sequence in this region. Interestingly, deletion of *mraW* abrogated the production of the frameshifted construct similar to the abrogation of production of *cmpA*^1–20^-*lacZ* (Fig. 4F). We therefore conclude that a property intrinsic to the *cmpA* nucleotide sequence, and not codon usage, is likely required for MraW regulation of *cmpA* translation.

**Figure 4.**
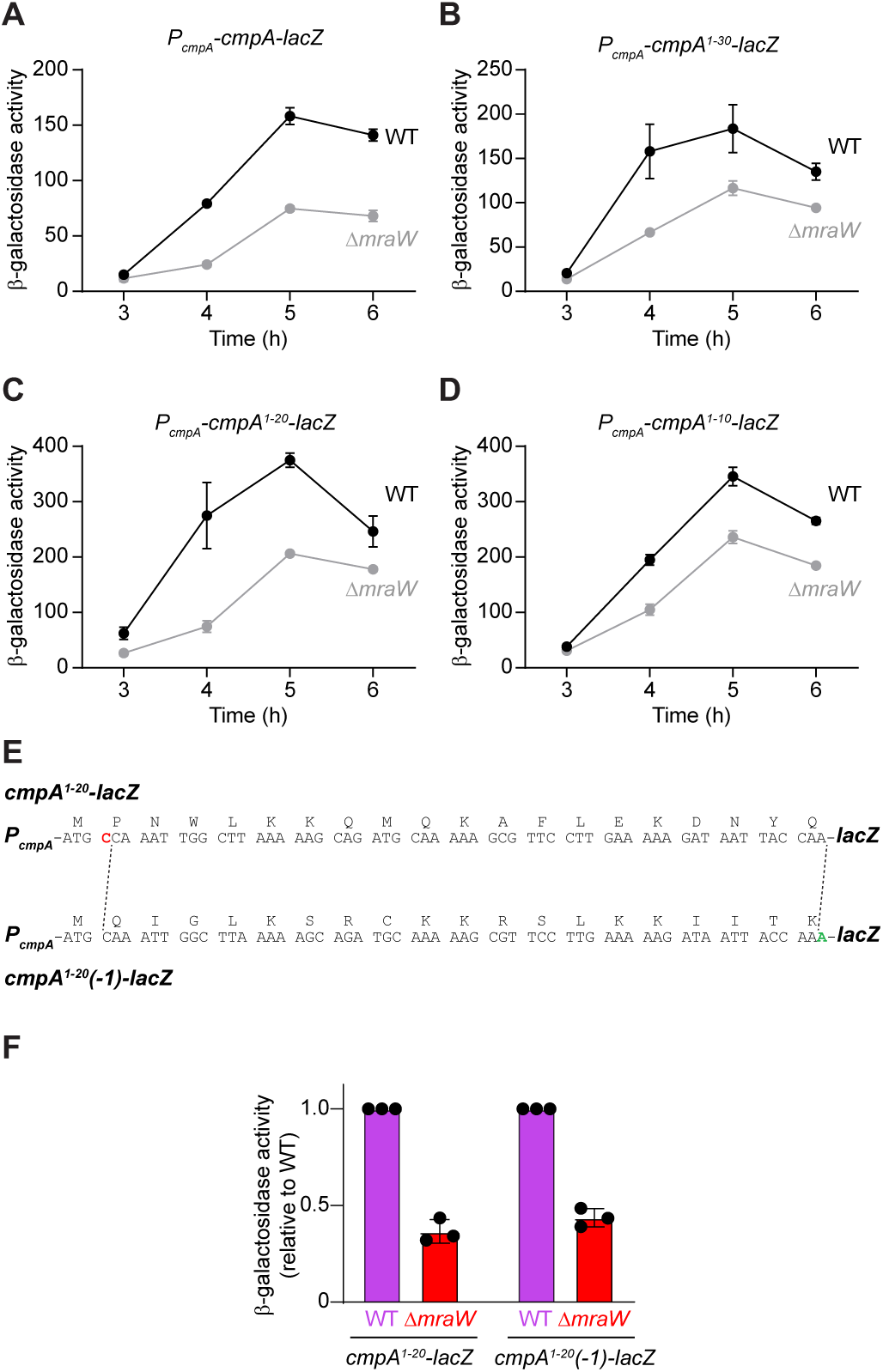
The nucleotide sequence of the first 20 codons of *cmpA* is required for MraW regulation. (A-D) β-galactosidase activity arising from translational fusions of (A) full length *cmpA*, (B) *cmpA^1-^*^30^, (C) *cmpA^1-20^*, or (D) *cmpA^1-10^* fused to *lacZ* in the (black) presence or (gray) absence of *mraW* at indicated time points after induction of sporulation. Data points represent mean (n = 3 biological replicates; errors: S.D.). Strains: SE230, ZP313, ZP441, ZP446, ZP439, ZP444, ZP437, ZP442. (E) Illustration of the sequence changes to create a -1-frame shift within the first 20 codons of the *cmpA* sequence. Top: *cmpA^1-20^* sequence; bottom: *cmpA^1-20^(-1)* sequence. Red indicates a deleted nucleotide in the first *cmpA* codon; green indicates an added nucleotide at codon 20 to restore the ORF with the *lacZ* fusion. Amino acids encoded by each codon in either construct displayed above the corresponding codons. (F) β-galactosidase activities relative to WT of *cmpA^1-20^-lacZ* or *cmpA^1-20^(-1)-lacZ* in (purple) presence or (red) absence of *mraW* measured 4 h after induction of sporulation. Strains: ZP439, ZP444, ZP463, and ZP469. Bars represent mean (n = 3 biological replicates); errors: S.D.

### A predicted early stem-loop structure in cmpA mRNA is required for MraW regulation of translation

RNA structure prediction models revealed that the nucleotide sequence between the ribosome binding site (RBS) and the AUG start codon (a sequence that we term the “intervening sequence”) along with the first 6 codons of *cmpA* forms a stem-loop structure (Fig. S4). The nucleotides contributing to this structure are relatively well conserved across *cmpA* orthologs (Fig. S5) and, with the exception of the *B. anthracis* ortholog, the different orthologs are predicted to form a stem-loop structure near the beginning of the ORF in which the intervening sequence participates at least partially in base pairs with early codons in the *cmpA* ORF (Fig. 5A-H).

**Figure 5.**
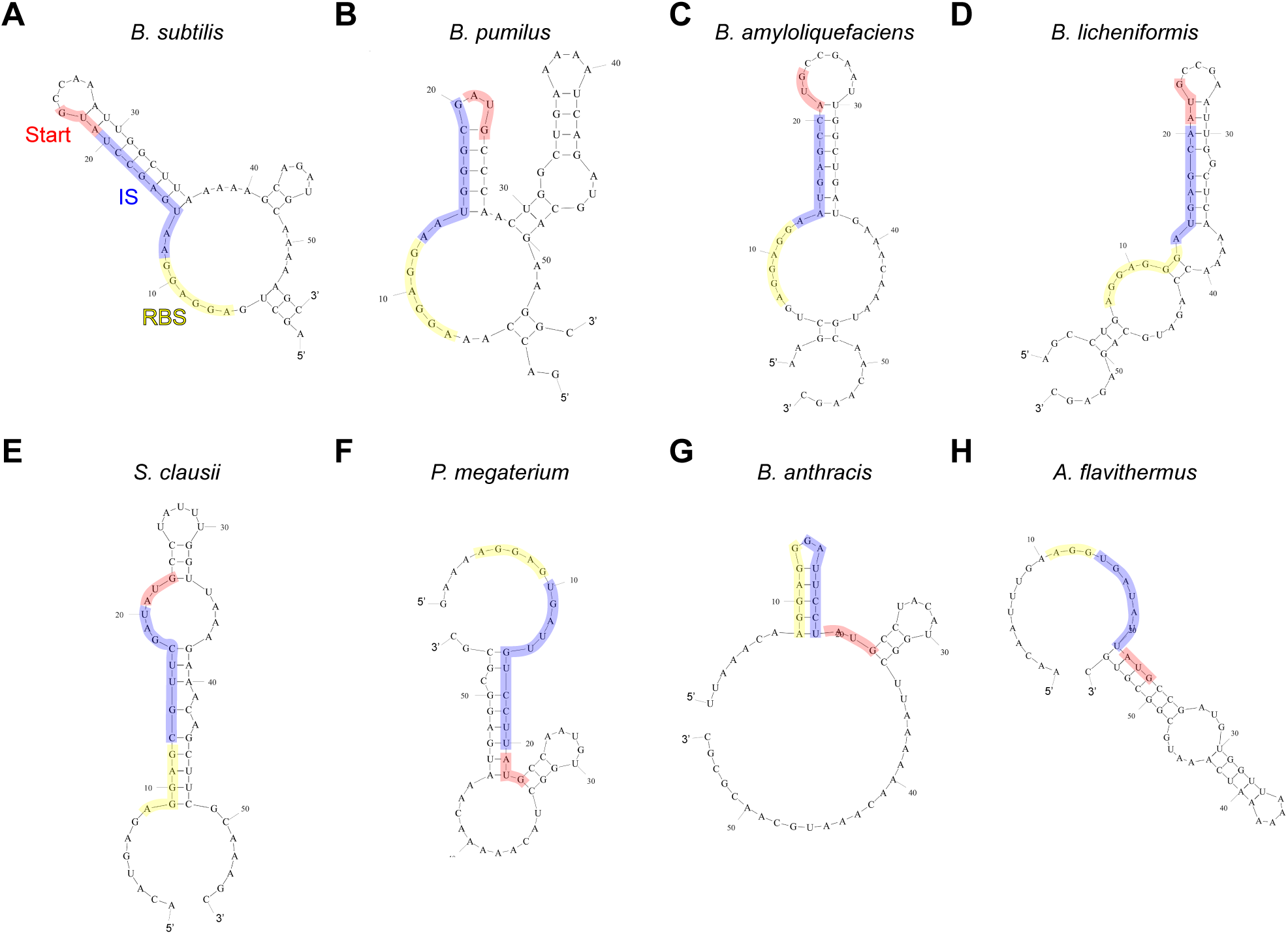
Conservation of structural elements in the 5’ region of *cmpA* mRNA. mRNA structure predictions of the 20 nucleotides upstream of start plus the first 35 nucleotides of the *cmpA* ORF generated by Mfold from (A) *B. subtilis*, (B) *Bacillus pumilus*, (C) *Bacillus amyloliquefaciens*, (D) *Bacillus licheniformis*, (E) *Shouchella clausii*, (F) *Priestia megaterium*, (G) *Bacillus anthracis*, and (H) *Anoxybacillus flavithermus*. RNA sequence alignment of this region is shown in Figure S4. RBS is highlighted in yellow; start codon: red; intervening sequence between the RBS and start codon (IS): blue.

To test if this structure is important for MraW-mediated regulation of *cmpA* translation, we constructed several mutants that disrupted or restored this structure and assessed the impacts of these changes on CmpA production in vitro and in vivo. First, we replaced the entire intervening sequence (5’-AAUGAGCCU) with 5’-AAAAAAU to disrupt the stem structure and measured the production of CmpA-GFP in vitro in a combined transcription-translation reaction. As seen previously (Fig. 2A) ribosomes isolated from Δ*mraW* cells produced ∼2-fold lower levels of CmpA-GFP compared to ribosomes isolated from WT cells (Fig. 6A). In addition, the production of CmpA-GFP was much lower than GFP production (Fig. 2A) alone even by WT ribosomes suggesting that the presence of the CmpA sequence is inhibitory to even WT ribosomes. However, in vitro translation of the mRNA harboring a disrupted stem resulted in an increase of CmpA-GFP production by Δ*mraW* ribosomes, similar to levels produced by WT ribosomes that translated the mRNA containing the stem (Fig. 6A). Interestingly, WT ribosomes displayed even more increased production of CmpA-GFP from the stem-disrupted transcript. To test if the increased production of CmpA from the mutant transcript produced higher quantities of CmpA that are physiologically relevant, we tested the effect of this construct in vivo during sporulation. Overexpression of *cmpA* from a constitutive promoter resulted in a sporulation defect that was corrected by deletion of *mraW* (Fig. 6B, lanes 2-3). Overexpression of the *cmpA* transcript containing the stem disruption also resulted in a similar sporulation defect (Fig. 6B, lane 4), but deletion of *mraW* did not correct this sporulation defect (Fig. 6B, lane 5). These findings suggest that the increase in CmpA-GFP production by *ΔmraW* ribosomes in vitro from a template in which the stem was disrupted translates to an observable phenotype in vivo.

**Figure 6.**
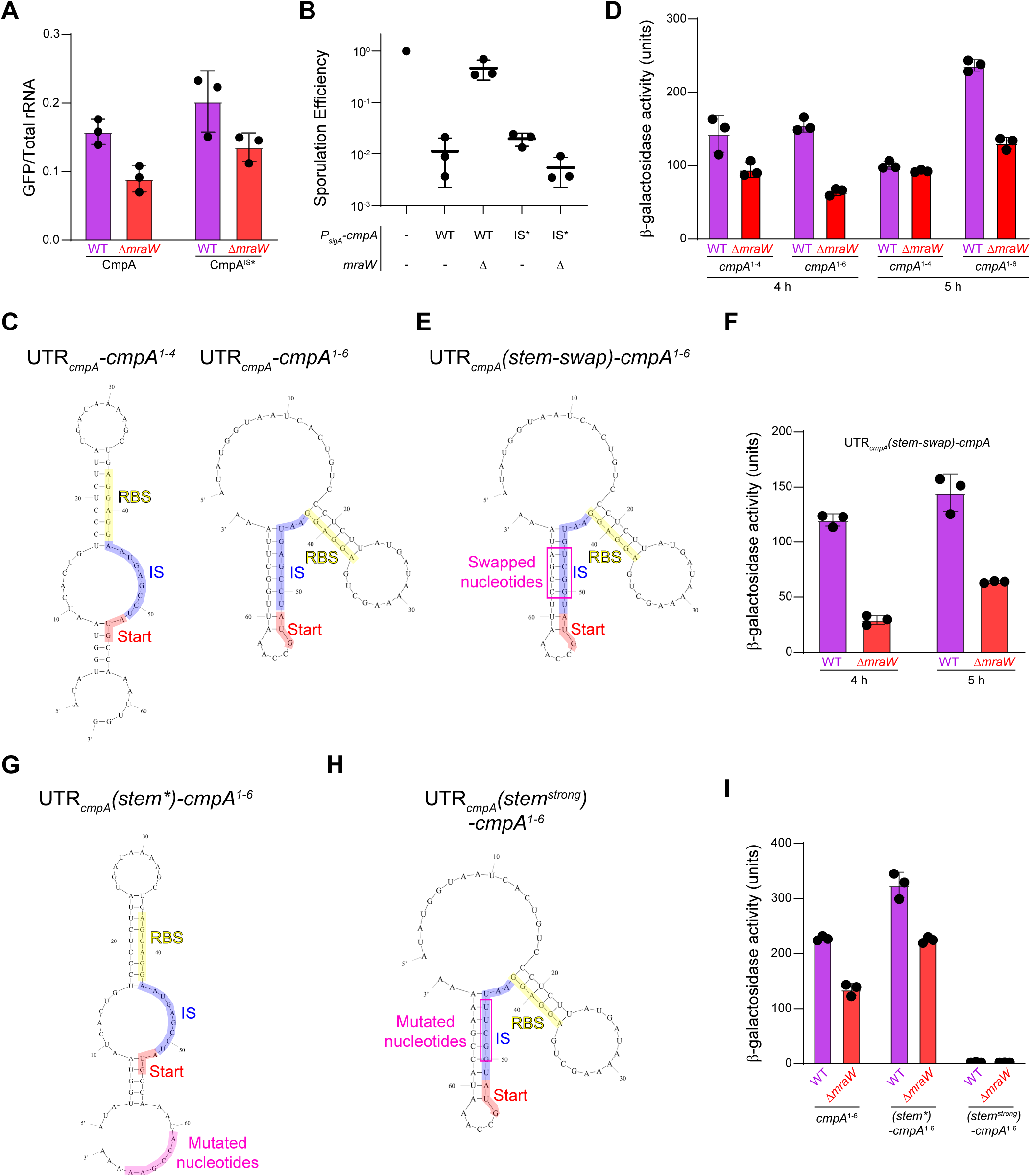
The stem-loop structure is required for MraW mediated translational regulation of *cmpA* mRNA. (A) CmpA-GFP fluorescence generated from an in vitro transcription/translation assay of a construct containing either the native *cmpA* IS sequence (“AAUGAGCCU”) or a mutant (IS*) sequence (“AAAAAAU”) using ribosomes purified from WT (purple) or Δ*mraW* (red) *B. subtilis* (strains PY79 or ZP1) after 3 hours of growth in LB 10/10/10 media. Bars represent mean (n = 3 independent trials using 3 separate sets of ribosomes purified from each strain); errors: S.D. (B) Sporulation efficiencies of strains in the presence of either wild-type (WT) or IS mutant (IS*) *cmpA* that is constitutively produced (*P_sigA_-cmpA*) in the presence or absence of *mraW*. Bars represent mean (n = 3 biological replicates); errors: S.D. Strains: PY79, IT686, ZP116, ZP560, and ZP562. (C) Mfold predictions of the 5’ UTR and either the first four or six codons of *cmpA*. (D) β-galactosidase activity arising from the translational fusion of the 5’UTR and either first four or six codons of *cmpA* to *lacZ* in the (purple) presence or (red) absence of *mraW* at indicated time points after induction of sporulation. Data points represent mean (n = 3 biological replicates; errors: S.D.). Strains: ZP451, ZP457, ZP473, and ZP481. (E) Mfold structure prediction in which several nucleotides comprising the stem were swapped. (F) β-galactosidase activity arising from the translational fusion of the swapped sequence in the presence (purple) and absence (red) of *mraW* at indicated time points after induction of sporulation. Data points represent mean (n = 3 biological replicates; errors: S.D.). Strains: ZP503 and ZP497. (G-H) Mfold predictions of the 5’ UTR and first six codons of *cmpA* with (G) codon 4 mutated to “ACC” and codon 5 mutated to “GAA” to disrupt the stem-loop structure, and (H) a stronger stem structure created by mutation of nucleotides in (G). (I) β-galactosidase activities from the constructs in (G-H) in the presence (purple) or absence (red) of *mraW* at 5 hours post-induction of sporulation. Data points represent mean (n = 3 biological replicates; errors: S.D.). Strains: ZP473, ZP481, ZP532, ZP534, ZP568, and ZP570.

Next, we sought to further probe the importance of the stem loop structure in vivo. Based on the structure prediction, the 5’ untranslated region and first 6 codons of *cmpA*, but not codons 1-4, are sufficient to form the stem loop (Fig. 6C). Consistent with this prediction, deleting *mraW* reduced synthesis of CmpA^1-6^-LacZ, but not CmpA^1-4^-LacZ (Fig. 6D). To test the sufficiency of stem formation for MraW regulation, we swapped 4 nucleotides in the intervening sequence with the corresponding 4 nucleotides in the *cmpA* coding region so that the predicted stem would not be disrupted. Deleting *mraW* resulted in reduced synthesis of this stem-swapped variant of CmpA-LacZ (Fig. 6F), consistent with the model that the presence of the early stem structure in the mRNA requires MraW-modified ribosomes for robust translation.

We next examined if disrupting the stem-loop structure in vivo would abrogate the need for MraW-modified ribosomes for efficiently translating *cmpA* mRNA. Using the CmpA^1-6^-LacZ template, we mutated codons 4 and 5 from “TGG CTT” to “ACC GAA” to disrupt the stem-loop structure (Fig. 6G). Translation of this mutant *cmpA* transcript (*cmpA^1-6^(stem*)-lacZ*) showed reduced dependence on the presence of MraW-modified ribosomes and resulted in similar levels of β-galactosidase activity as the native CmpA^1-6^-LacZ produced by wild-type ribosomes (Fig. 6I). Interestingly, similar to what we observed in vitro (Fig. 6A), disrupting the stem-loop also increased expression by wild-type ribosomes in vivo, again suggesting that the presence of the stem-loop limits CmpA production even in wild-type cells (Fig. 6I). Finally, we noticed that the native stem structure in *cmpA^1-6^-lacZ* contained two instances of imperfect or no base pairing (either U-G or U-U). We therefore introduced additional mutations to change the U-U non-base pair to A-U and the U-G base pair to A-U to construct *cmpA^1-6^(stem^strong^)-lacZ* (Fig. 6H). Surprisingly, the resulting stable stem structure completely blocked translation, even in otherwise WT cells (Fig. 6I). The data are therefore consistent with a model in which *cmpA* can be more efficiently translated by MraW-modified ribosomes due to the enhanced ability of these ribosomes to engage mRNAs with secondary structures that occlude early codons in the ORF.

### Translation of other mRNAs with predicted stem-loops are impacted in ΔmraW

We next examined the proteome of *B. subtilis*, either WT or cells harboring a mutant allele of *spoIVA*, during vegetative growth and sporulation using mass spectrometry to assess the impact of MraW on the production of other proteins besides CmpA. The raw data revealed scores of proteins that were differentially regulated in the presence and absence of MraW (Supplemental Dataset 1). Of note, loss of MraW led to wild type levels of SpoIVA^T*^ consistent with our model of lower CmpA levels in Δ*mraW* inhibiting mutant SpoIVA degradation (Fig. S6). To identify proteins whose levels were altered directly because of MraW, we employed three criteria to narrow the list of impacted proteins, based on the assumption that the loss of MraW should impact translation of a given transcript to a similar extent regardless of the allele of *spoIVA* the strain harbored. First, we considered only target proteins whose production changed at least 2-fold; second, the protein had to be detected in all analyzed strain backgrounds; third, the change in production had to be reproducible and statistically significant across three replicates (*P* < 0.05).

Using these relatively stringent criteria, we identified two proteins that we further analyzed: Isp, a protease involved in the transition to stationary phase ^50^; and YukJ, a protein of unknown function that is likely upregulated in stationary phase ^51^. Like the *cmpA* transcript, the 5’ end of the *isp* mRNA is predicted to harbor a stem loop structure, but unlike *cmpA* (whose stem occludes the untranslated intervening sequence and several early codons) the *isp* stem loop exclusively comprises sequences upstream of the RBS, the intervening sequence, and just the start codon (Fig. 7A). Fusion of just the *isp* promoter and this 5’ untranslated region of *isp* to *lacZ* resulted in the MraW-dependent production of β-galactosidase at 0h, 1.5 h, and 4 h after induction of sporulation (Fig. 7B). This dependence on MraW persisted when we swapped the nucleotides in the predicted stem (Fig. 7C-D), suggesting that the stem loop structure in the 5’ UTR of *isp* inherently reduces translation efficiency of this transcript and that MraW-modified ribosomes are capable of efficiently translating this mRNA. Consistent with this model, disrupting the stem loop by exchanging the intervening sequence (normally GCTTTTTT) with the short intervening sequence that we used in the *cmpA* experiments (AAAAAAT) (Fig. 7E) resulted in enhanced translation in Δ*mraW* cells and, like what we observed with the similarly mutated *cmpA* transcript, also enhanced expression in otherwise WT cells (Fig. 7F). Conversely, altering the stem so that it became more stable by substituting the A-T base pairs to G-C resulted in drastically reduced expression in both WT and Δ*mraW* cells (Fig. 7G-H), similar to what we observed when we enhanced the stability of the *cmpA* stem (Fig. 6I).

**Figure 7.**
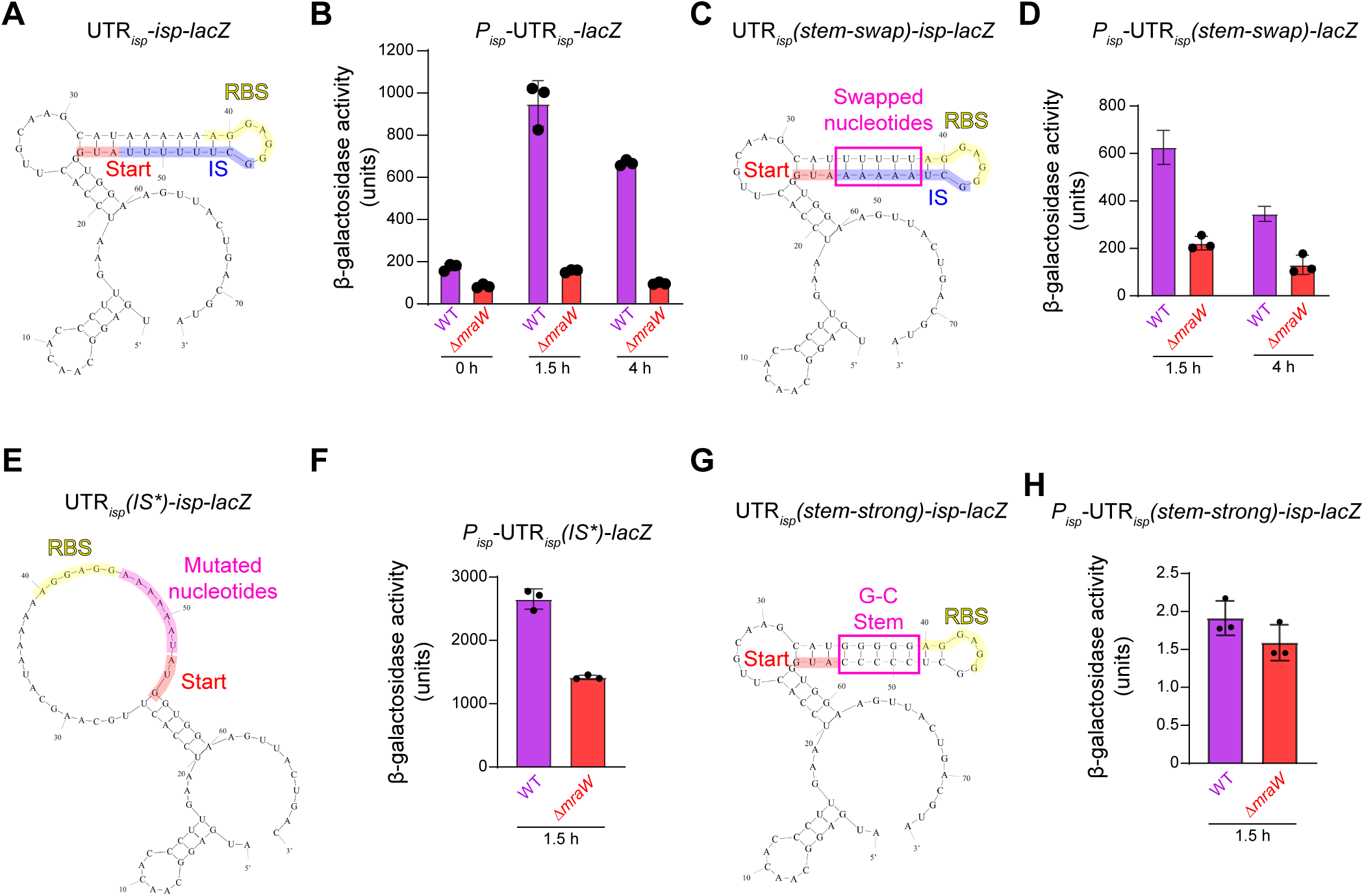
MraW is required for the efficient translation of other transcripts with early stem-loop structures. (A) Mfold predicted structure of the 5’ UTR and start codon of *isp* with six codons from *lacZ*. (B) β-galactosidase activity arising from the fusion of the promoter and 5’UTR of *isp* to lacZ at indicated time points after induction of sporulation in the presence (purple) and absence (red) of MraW. Data points represent mean (n = 3 biological replicates; errors: S.D.). Strains: ZP507 and ZP513. (C) Mfold predicted structure of the 5’UTR and start codon of isp with six codons from lacZ in which the nucleotides that form the stem structure have been swapped with each other. (D) β-galactosidase activity arising from the stem-swapped mutant of *isp* at indicated time points after induction of sporulation in the presence (purple) and absence (red) of MraW. Data points represent mean (n = 3 biological replicates; errors: S.D.). Strains ZP509 and ZP515. (E) Mfold predicted structure of the 5’UTR and start codon of isp in which the IS has been mutated from “GCUUUUUU” to “AAAAAAU” fused to lacZ. (F) β-galactosidase activity arising from the IS mutant of *isp* at 1.5 hours after induction of sporulation in the presence (purple) and absence (red) of MraW. Data points represent mean (n = 3 biological replicates; errors: S.D.). Strains ZP577 and ZP583. (G) Mfold predicted structure of the 5’ UTR and start codon of *isp* in which the A-T base pairs were substituted with G-C base pairs, fused to *lacZ*. (H) β-galactosidase activity arising from the G-C mutant of *isp* at 1.5 h after induction of sporulation in the presence (purple) and absence (red) of MraW. Data points represent mean (n = 3 biological replicates; errors: S.D.). Strains ZP511 and ZP517.

The *yukJ* transcript is also predicted to form an early stem loop structure, but here, the structure comprises exclusively codons 2-11 (and partially codon 12) and excludes the start codon and any upstream untranslated sequences (Fig. S7A). YukJ protein levels were reduced in cells lacking MraW at each time point analyzed (Fig. S7B), but fusing the *yukJ* promoter alone to *lacZ* showed no MraW-dependent effect, indicating that the impact of Δ*mraW* on YukJ expression is occurring at the translation step (Fig. S7C). We conclude that the regulation of translation via secondary structures that limit translation efficiency, which evolved in select transcripts, is a general method in *B. subtilis* by which precise cellular concentrations of certain proteins can be maintained, and that this regulation is dependent on ribosomes whose 16S rRNA has been methylated by MraW.

## DISCUSSION

In this study, a selection for spontaneous suppressor mutations that correct the sporulation defect of cells producing a defective variant of the spore coat protein SpoIVA revealed a potential role for rRNA modifications in regulating the proper completion of the sporulation developmental program. A loss of function mutation in the widely conserved 16S rRNA methyltransferase, MraW, enhanced sporulation efficiency even in the presence of defective SpoIVA alleles, suggesting that these suppressor mutants bypassed the coat assembly checkpoint ^44^. Using a candidate approach, we determined that the transcript encoding the negative sporulation regulator CmpA was poorly translated in the absence of MraW, thereby suggesting a model by which cells harboring mistakes in coat assembly could nonetheless proceed through the sporulation program. We discovered that the *cmpA* transcript harbors a stem-loop structure that occludes several early codons in the *cmpA* ORF, resulting in a transcript that requires MraW-modified ribosomes to fine-tune the translation (likely translation initiation) of the transcript encoding this negative regulator: not so robustly as to inhibit sporulation in normal cells, but simultaneously producing enough CmpA to arrest sporulation if a coat assembly mistake arises during sporulation. A high throughput analysis revealed other transcripts harboring similar stem-loop structures near the 5’ end of the ORF that also regulate translation of these mRNAs in an MraW-dependent manner. We propose that specific transcripts, perhaps especially those that encode proteins whose cellular levels should be limited (like CmpA), exploit this rRNA modification to regulate protein production to ensure the faithful progression of bacterial development. Evolution of these mechanisms may have been facilitated by the uncoupling of transcription and translation observed in *B. subtilis* and predicted for other species ^52^. While a similar stem-loop mediated mechanism could also be present in *E. coli* for example, the tight coupling of transcription and translation in this organism ^53,54^ may preclude a mechanism by which structural elements in the mRNA could fold before the engagement with the ribosome.

Consistent with our model of secondary structure-mediated translational regulation of select transcripts, even though deletion of *mraW* does not result in an obvious growth defect, we observed in vitro that purified ribosomes not modified by MraW displayed an inherently lower efficiency in translating even optimized transcripts, and that this impairment was magnified in *cmpA* transcripts that harbor the translation-inhibitory stem-loop structure. We therefore speculate that the requirement for MraW may be more evident when cells are exposed to growth conditions that may stabilize translation-inhibitory structural elements present in select mRNAs. Interestingly, a recent report showed that *B. subtilis* lacking MraW exhibits defective growth at cold temperatures (18 °C) and in recovery from cold shock ^55^. Δ*mraW* cells did not display obvious defects in ribosome assembly after cold shock, but instead a reduction in the presence of polysomes, leading us to speculate that this defect is perhaps due to the stabilization of translation-inhibitory mRNA structural elements at reduced temperatures. The deletion of *mraW* resulted in a less severe defect in *E. coli* ^55^, indicating that this level of MraW-mediated regulation is species-specific.

The transcript specificity for the requirement of MraW was evidenced by the relative scarcity of proteins whose production was reproducibly affected in vivo in the absence of MraW (consistent again with the lack of an obvious growth phenotype of an *mraW* deletion mutant). RNA secondary structure is a common mechanism for the fine-tuning of post-transcriptional regulation, with repressive effects of the structured regions often relieved by small RNAs, riboswitches, and RNA thermometers ^56–62^. We propose that the presence of such secondary structures in the ORFs of select *B. subtilis* mRNAs represents a different regulatory mechanism that perhaps requires rRNA modifications for efficient engagement of the trapped codons to facilitate efficient translation initiation in cases where the start codon is occluded, and to facilitate efficient translation elongation in cases where the stem-loop forms immediately after the start codon, as exemplified in the *yukJ* transcript. Once the stable interaction between the modified rRNA and mRNA codons are formed, the intrinsic helicase activity of ribosomes ^63^ is likely more efficient at unwinding the mRNA. Indeed, structural data indicate that m4mC1402 directly engages with the mRNA backbone and forms key chemical interactions to stabilize the P-site of the small ribosomal subunit ^5,20^. This role in increased translation efficiency for C1402 methylation is distinct from, but complementary to, another phenotype that was previously described for an *mraW* deletion mutant wherein translation from GUG and UUG start codons was impaired, coupled with mis-initiation at AUU codons ^20^. Combined with the observations we report here, we hypothesize that loss of MraW impairs the ability of the small subunit to efficiently engage start codons and/or codons immediately following the start codon that are present in the context of these structured regions, which could explain why CmpA production is not completely abrogated in the Δ*mraW* mutant and why there are not severe growth defects in Δ*mraW* strains. In sum, given the widespread conservation of *mraW* in bacterial genomes ^21^, combined with the presence of orthologous methyltransferases and rRNA modifications in chloroplasts and mitochondria ^64,65^, it appears that rRNA modifications may broadly play a key role in accurate start site selection by the ribosome, and that secondary structures in the mRNA that encode sensitive regulatory proteins may be a general strategy to regulate the production of those factors.

## ACKNOWLEDGEMENTS

We thank S. Gottesman, G. Storz, A. Khare, and A. Buskirk for discussions, members of the K.S.R. lab for comments on the manuscript, and M. Wong of the CCR Genomics Core for the TapeStation analysis. This research was supported by the Intramural Research Program of the National Institutes of Health (NIH), National Cancer Institute, Center for Cancer Research (K.S.R.). The contributions of the NIH authors are considered Works of the United States Government. The findings and conclusions presented in this paper are those of the authors and do not necessarily reflect the views of the NIH or the U.S. Department of Health and Human Services.

## DECLARATION OF INTERESETS

The authors declare no competing interests.

## DATA AVAILABILITY

The mass spectrometry proteomics data have been deposited to the ProteomeXchange Consortium via the MassIVE partner repository with the dataset identifier PXD074358.

## EXPERIMENTAL PROCEDURES

### Strain Construction

All strains used in this study were derived from the parental *B. subtilis* strain PY79 ^66^. The Δ*mraW*::*erm* deletion was obtained from an ordered deletion library ^67^ whose chromosomal DNA was transformed into PY79 to generate strain ZP1. Insertion of *spoIVA* at the *sacA* locus was achieved by cloning the *spoIVA* CDS plus 450 bp upstream and 360 bp downstream into RL4165 before transformation into *B. subtilis*. SpoIVA variants were derived from that plasmid (pZP30) using QuikChange mutagenesis. Fusions to *lacZ* were generated using either synthesized gene fragments (IDT) or QuikChange mutagenesis of previously constructed plasmids derived from RL5037 for insertion of the fusions at the *amyE* locus. The constitutively expressed *cmpA-GFP* fusion was constructed by using QuikChange mutagenesis to insert two early in-frame stop codons into the *lacI* gene in pSE24 ^68^.

### General Methods

SpoIVA, MraW, and σ^A^ levels were measured using immunoblotting of sporulating cell extracts prepared as previously described ^43^. All proteins were detected using rabbit antiserum raised against recombinant SpoIVA (1:10,000 dilution) ^69^, MraW (1:10:000 dilution), and σ^A^ (1:5000 dilution) ^70^ (Covance) as the primary antibody and goat StarBright Blue 700 (1:10,000 dilution) (Bio-Rad) as the secondary antibody. Immunoblots were imaged on a ChemiDoc^TM^ MP Imaging System (BioRad). GFP production was quantified from 100 µl cultures of sporulating cultures at the indicated time points grown in a 96-well plate using a Synergy H1 microplate reader (BioTek) (488 nm excitation, 530 nm emission). For growth curves, overnight cultures were grown in lysogeny broth (LB) at 22 °C and diluted (1:100) into the indicated media in a 96-well plate and grown at 37 °C with shaking in a Synergy H1 microplate reader (BioTek) plate reader with O.D._600nm_ readings taken every 15 minutes.

### Sporulation Efficiency

Sporulation efficiencies were calculated by heating cultures at 80 °C for 20 min after 24 h of growth in Difco Sporulation Media at 37 °C (DSM) to remove non-sporulating or defective sporulating cells from the population. Serial dilutions of the heat killed cultures were then plated on LB to determine the number of colony forming units (cfu); sporulation efficiency is reported relative to heat-resistant cfu calculated for a parallel culture of WT (PY79).

### SpoIVA purification and ATP Hydrolysis Assay

Plasmid expressing *his_6_*-*spoIVA*^L59P^ was generated by QuikChange mutagenesis of pKR145 ^40^. His_6_-SpoIVA^L59P^ was purified as described previously ^71^. Purified protein (0.3 µM) was then incubated with varying concentrations of ATP at 37 °C for 1 h. The reactions were stopped by addition of 50 µl of 0.02 M EDTA. Freed phosphate from SpoIVA hydrolysis was measured using the colorimetric GTPase Assay Kit (abcam). A standard curve generated from known [P_i_] solution was used to determine the levels of free phosphate generated from SpoIVA hydrolysis ^42^. Reactions were measured at A_620nm_ and the data was fitted to a Michaelis-Menten curve using Prism software (GraphPad).

### Epifluorescence Microscopy

Cells expressing GFP-SpoIVA were induced to sporulate using the resuspension method ^72^. Samples were taken at 2.5 h after induction of sporulation, collected by centrifugation, and resuspended in a 5 µg ml^-1^ solution of FM4-64 in water to visualize membranes. Stained samples were placed on a glass bottom culture dish and covered with a 1% agarose pad. Cells were visualized with a DeltaVision Core microsocpe (Applied Precision/GE Healthcare) as described previously ^73^, and images were processed with SoftWorx and ImageJ software.

### Ribosome Purification

*B. subtilis* ribosomes were purified as previously described ^65,74^. Briefly, overnight cultures of WT (PY79) or *ΔmraW* (ZP1) grown at 22 °C in LB medium (made using the 10/10/10 formulation) were diluted into 500 ml of fresh LB. Cells were harvested after 3 h of growth at 37 °C in lysogeny broth (made using the 10/10/10 formulation) and cell pellets were stored at -80 °C. Cells were then resuspended in 2 ml of lysis buffer (20 mM Tris pH 7.6, 10 mM magnesium acetate, 150 mM potassium acetate, 0.1% Triton-X-100, 1 mg ml^-1^ lysozyme, 100 U ml^-1^ DNase I) and incubated at 30 °C for 30 min. Lysed samples were then cleared by centrifugation at ∼8,500 × *g* (14,000 RPM in an Eppendorf MiniSpin Plus rotor) for 5 min. ∼500 µl of supernatant were gently layered on a 500 µl sucrose cushion (50 mM Tris-acetate at pH 7.5, 1 M potassium acetate, 15 mM magnesium acetate, 1.44 M sucrose, 1 mM dithiothreitol (DTT), 0.1% Complete Protease Inhibitor) in 11 × 34 mm PC tube (Beckman) and centrifuged for 1 h at ∼430,000 × *g* (100,000 rpm in a Beckman TLA 120.2 rotor). The ribosome-containing pellet was resuspended in 50 µl Buffer B (50 mM Tris-acetate at pH 7.5, 50 mM potassium acetate, 5 mM magnesium acetate, 1 mM DTT). Ribosome samples were analyzed using a Nanodrop spectrophotometer to measure RNA concentration, by separation on a non-denaturing agarose gel to assess rRNA integrity, and by Coomassie-PAGE to assess relative abundance of ribosomal proteins. Quality of purified ribosomes was further analyzed using the TapeStation system (Agilent) to determine RIN values. Purified ribosomes were used immediately for the in vitro transcription and translation assay (described below) and stored at -80 °C.

### In vitro Transcription and Translation

In vitro transcription/translation was performed using the PURExpress In Vitro Protein Synthesis kit (NEB) using ribosomes purified from WT or *ΔmraW* cells. DNA templates for the reaction were generated using PCR with primers designed using kit guidelines and including sequences from pSE24 to amplify the GFP sequence. Reactions (25 µl) were run in biological triplicate largely following the manufacturer’s protocol using 4 µl of the purified ribosome preparation and the addition of 1 µl of RNase Out ribonuclease inhibitor (Invitrogen). Reactions were incubated at 37 °C for 4 h and were stopped by incubation on ice. Reactions were then mixed with 25 µl of sterile diH_2_O and fluorescence from the synthesized GFP was measured using a Synergy H1 microplate reader (BioTek) (488 nm excitation, 530 nm emission).

### β-galactosidase assays

Cells expressing *lacZ* fusions were induced to sporulate using the resuspension method. At indicated time points, the O.D._600nm_ of the culture was measured and 1 ml of culture was harvested and stored at -80 °C. Assays were conducted largely as described ^68^. Briefly, harvested cells were resuspended in 1 ml of Z-buffer (0.06M Na_2_HPO_4_-7H_2_O, 0.04M Na_2_HPO_4_-H_2_O, 0.01M KCl, 0.001M MgSO_4_, 0.05M β-mercaptoethanol) and 50 µl of each sample were aliquoted into a well of a sterile 96-well plate. These samples were diluted with 50 µl of Z-buffer with 0.4 mg ml^-1^ lysozyme and incubated at 37 °C for 30 min. After incubation, 20 µl of 4 mg ml^-1^ o-Nitrophenyl β-D-galactopyranoside in Z-buffer were added to each sample and the absorbance at 420 nm was measured every minute for one hour using a Synergy H1 microplate reader (BioTek) plate reader. Slopes were determined using Excel and arbitrary units were calculated.

### mRNA folding

mRNA structure predictions were conducted using the Mfold RNA secondary structure prediction program ^75,76^ and verified with Vienna Fold ^77^. Sequence identities of the 5’ UTR were determined using the SubtiWiki genome browser ^78^, which identifies upshifts indicating the transcription start site. The standard parameters were used for both modeling platforms.

### Sequence Conservation

The DNA sequences from 100 nucleotides upstream to the first 10 codons of the *cmpA* genes in various spore forming bacteria^1^ were obtained from NCBI Sequence Viewer 3.35.1 ^79^. Sequences were aligned with Clustal Omega 1.2.4 ^80^ first, then gaps were adjusted manually to improve the alignment in JalView 2.11.4.1 ^81^.

### Mass spectrometry

*B. subtilis* cell lysis was performed as follows. Frozen cell pellets harvested from 20 ml of sporulating cultures were suspended in 250 µl modified RIPA buffer (150 mM NaCl, 50 mM Tris-HCl at pH7.4, 1% NP-40, 0.25% Sodium deoxycholate, and 1 mM EDTA) containing protease inhibitors (Roche). To facilitate lysis, 25 µg lysozyme were added to each sample, and the tubes were incubated at 37 °C for 30 min, shaking at 750 rpm. Samples were mixed 1:1 with urea lysis buffer (6 M Urea, 25 mM HEPES at pH 8.0, 5 mM EDTA) containing the protease inhibitors and lysed using a TissueLyser bead mill homogenizer (Qiagen) for 2 min, repeated three times to ensure complete lysis. Samples were then centrifuged at 12000 × *g* for 15 min at 10 °C. After centrifugation, supernatants were transferred and protein concentrations measured using the BCA method (Thermo Scientific). Next, 100 µg of each protein sample was adjusted to a volume of 50 µl with urea lysis buffer, mixed with an equal volume of 50 mM TEAB, and reduced with 10 mM TCEP at 56 °C for 45 min in a shaker (750 rpm). Samples were then cooled and alkylated with 20 mM iodoacetamide at room temperature in the dark. Samples were further diluted with 200 µl 50 mM TEAB to reduce the urea concentration to below 2 M and proteins were digested with trypsin (1:50) and Lys-C (1:125) at 37 °C overnight. Following digestion, samples were acidified to a final concentration of 0.5% trifluoroacetic acid and desalted on a C18 spin column (Thermo Scientific).

Mass spectrometry analysis was performed using a Vanquish Neo UHPLC system (Thermo Fisher Scientific) coupled to an Orbitrap Exploris 480 mass spectrometer (Thermo Fisher Scientific) operating in data-independent acquisition (DIA) mode. Peptides were separated on a 75 μm × 50 cm EASY-Spray™ Neo UHPLC column (Thermo Fisher Scientific). The mobile phases consisted of (A) 0.1% formic acid (FA); (B) 0.1% FA, 80% acetonitrile. Peptides were separated using a 130 min gradient at a constant flow rate of 300 nl min^-1^: 2% B for 1 min, 2–32% B for 121 min, 32–90% B for 20 min. The Orbitrap Exploris 480 mass spectrometer was operated in positive ion mode with full-scan precursor spectra (MS1) acquired at a resolution of 120,000 over the m/z scan range of 420–680. The AGC target was set to 300% (3e6 ions absolute) with a maximum injection time of 100 ms. Fragment ion scans (DIA MS2) were collected using 60 isolation windows with a 4 m/z window width, covering a precursor range of 430–670 m/z. DIA windows were placed using the instrument’s automatic window-placement optimization. Fragment spectra were recorded in the Orbitrap at a resolution of 90,000, using stepped higher-energy collisional dissociation (HCD) with 25%, 27.5%, and 30% normalized collision energies. The DIA AGC target was set to 3000% (3e6 ions absolute) and the injection time was automatically determined. The scan range for fragment ions was 200–1800 m/z with DIA cycle time of 3 seconds.

DIA raw files were analyzed using DIA-NN 2.2.0 using an in-silico DIA-NN predicted spectral library from the UniProt *Bacillus subtilis* database (downloaded in 2025, 4191 sequences), allowing for cysteine carbamidomethylation and N-terminal methionine excision and 1 missed cleavage. The DIA-NN search included the following settings: Protein inference = Genes, Neural network classifier = Single-pass mode, Quantification strategy = Robust LC (high precision), Cross-run normalization = RT-dependent, Library Generation = IDs, RT and IM Profiling and Speed and RAM usage = Optimal results. Mass accuracy and MS1 accuracy were set to 0 for automatic inference. No share spectra, Heuristic protein inference and MBR were checked.

## SUPPLEMENTAL INFORMATION

**Figure S1.**
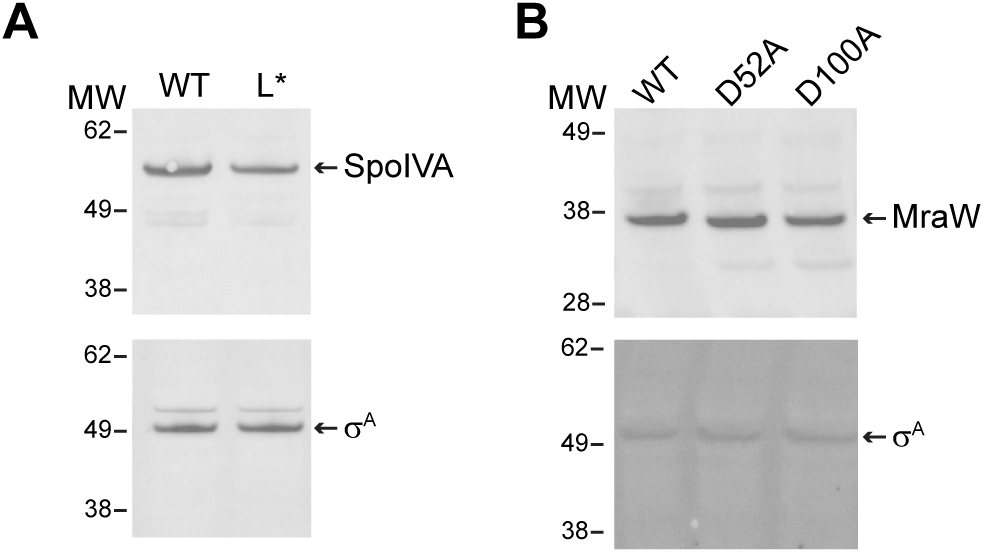
SpoIVA^L59P^ and MraW* are expressed at levels comparable to wild type versions of the proteins. (A) Cell extracts of sporulating *B. subtilis* producing either WT SpoIVA or SpoIVA^L^* harvested 2.5 h after induction of sporulation examined by immunoblotting using antisera to detect (top) SpoIVA or (bottom) SigA used as a loading control. Strains: KR394 and ZP47. (B) Cell extracts of sporulating *B. subtilis* producing either WT MraW, MraW^D52A^, or MraW^D100A^ harvested 4 h after induction of sporulation examined by immunoblotting using antisera to detect (top) MraW or (bottom) SigA. Strains: ZP134, ZP231, and ZP233.

**Figure S2.**
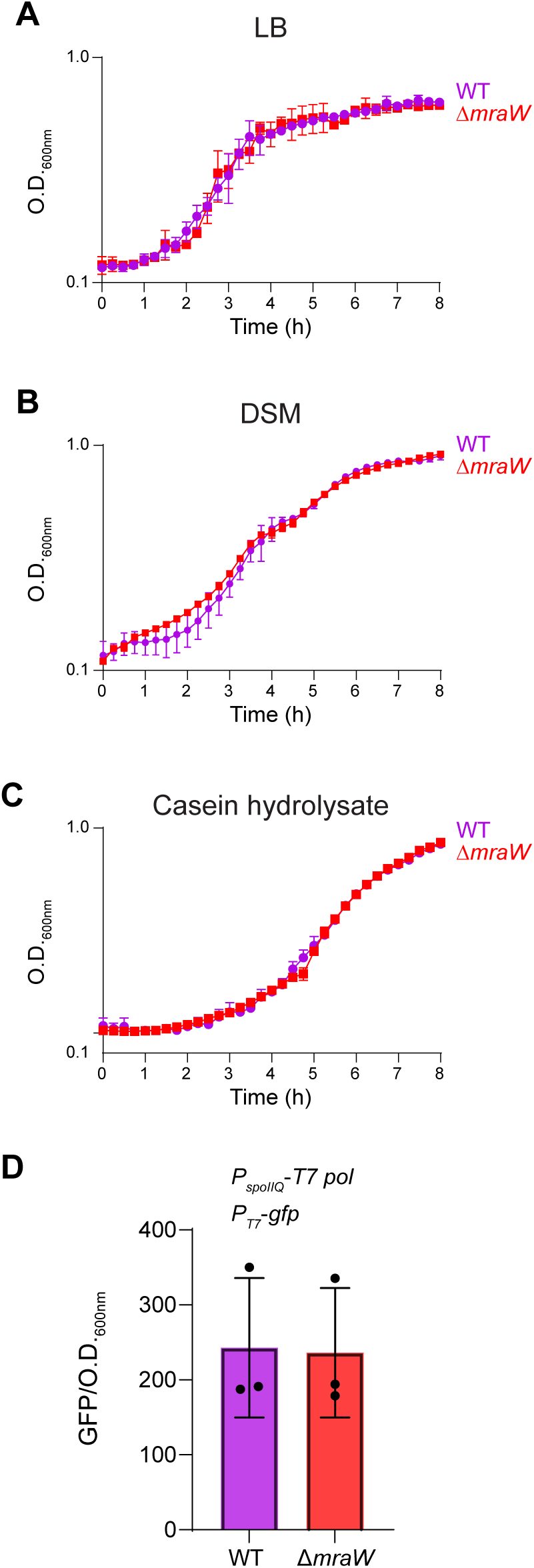
Growth rate of *B. subtilis* in several growth media is unaffected by deletion of *mraW*. Growth assayed by optical density measured at 600 nm (O.D._600nm_) at indicated time points of WT (purple) or Δ*mraW* cells grown in (A) lysogeny broth (LB), (B) Difco sporulation medium (DSM), or (C) casein hydrolysate medium. Data points represent mean (n = 3 independent cultures); errors: S.D. (D) Production of GFP in vivo whose gene expression was driven from a T7 promoter with a 5’ UTR that was identical to the template used in the in vitro transcription and translation assay in Figure 1A. Samples were analyzed after 18 hours of growth in LB 10/10/10 to allow for activation of the σF-dependent *spoIIQ* promoter that was driving T7 polymerase expression. Data points represent mean (n = 3 independent cultures); errors: S.D. Strains ZP572 and ZP574.

**Figure S3.**
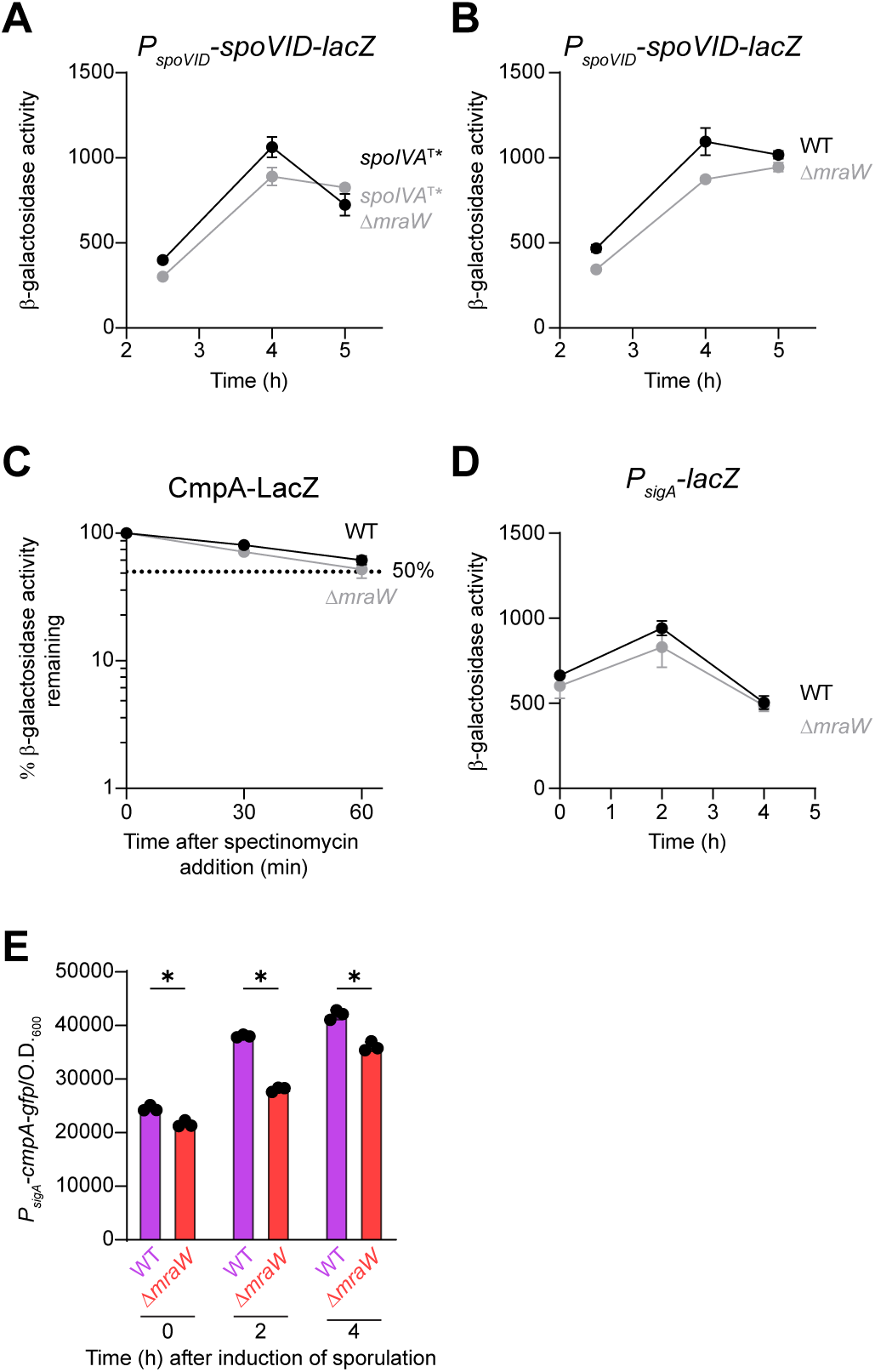
SpoVID production, CmpA turn-over rate, and *lacZ* expression, and are not impacted in Δ*mraW*. (A-B) β-galactosidase activity arising from translational fusion of the *spoVID* ORF to *lacZ* expressed from the *spoVID* promoter at indicated time points after induction of sporulation in the (black) presence or (gray) absence of *mraW* in cells harboring (A) *spoIVA*^T*^ or (B) WT *spoIVA*. Strains ZP405, ZP410, ZP339, ZP345. (C) β-galactosidase activity arising from translational fusion of *cmpA* ORF to *lacZ* after addition of 200μg mL-1 spectinomycin added 5.5 h after induction of sporulation to arrest translation in the (black) presence or (gray) absence of *mraW*. Activity is reported as % β-galactosidase activity remaining relative to t = 0. Data points represent mean (n = 3 independent cultures); errors: S.D. Strains SE230 and ZP313. (D) β-galactosidase activity arising from a transcriptional fusion of a constitutive promoter regulated by σ^A^ to *lacZ* ORF at indicated time points after induction of sporulation in the (black) presence or (gray) absence of *mraW*. Strains ZP363 and ZP361. Bars represent mean (n = 3 biological replicates); errors: S.D. (E) Levels of fluorescence generated by a CmpA-GFP fusion produced from a constitutive promoter regulated by σA relative to culture density (O.D._600nm_) at indicated time points after induction of sporulation in the (purple) presence or (red) absence of mraW. Statistical analysis: unpaired t-tests with the Bonferroni-Dunn correction for multiple comparisons, * indicates P-value < 0.05. Bars represent mean (n = 3 biological replicates); errors: S.D. Strains: ZP523 and ZP525.

**Figure S4.**
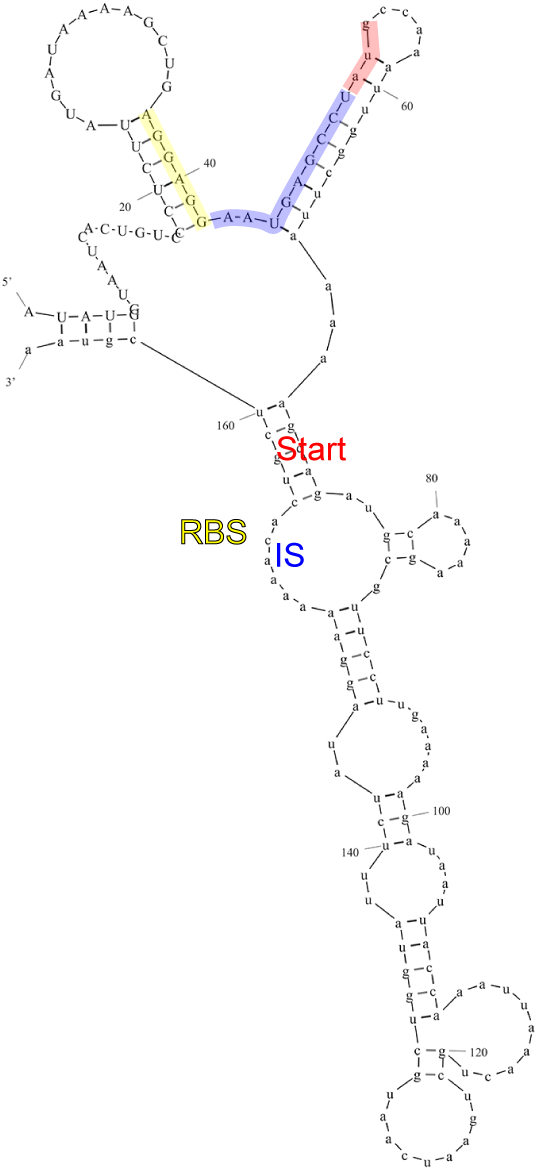
A stem-loop structure is predicted to occlude the start codon of *cmpA* mRNAs. Mfold prediction of the full length *cmpA* mRNA. The RBS is highlighted in yellow, the intervening sequence (IS) between the RBS and start is highlighted in blue, and the start codon (AUG) is highlighted in red. Sequence in uppercase represents the 5’UTR of *cmpA* mRNA and lowercase represents the *cmpA* coding sequence.

**Figure S5.**
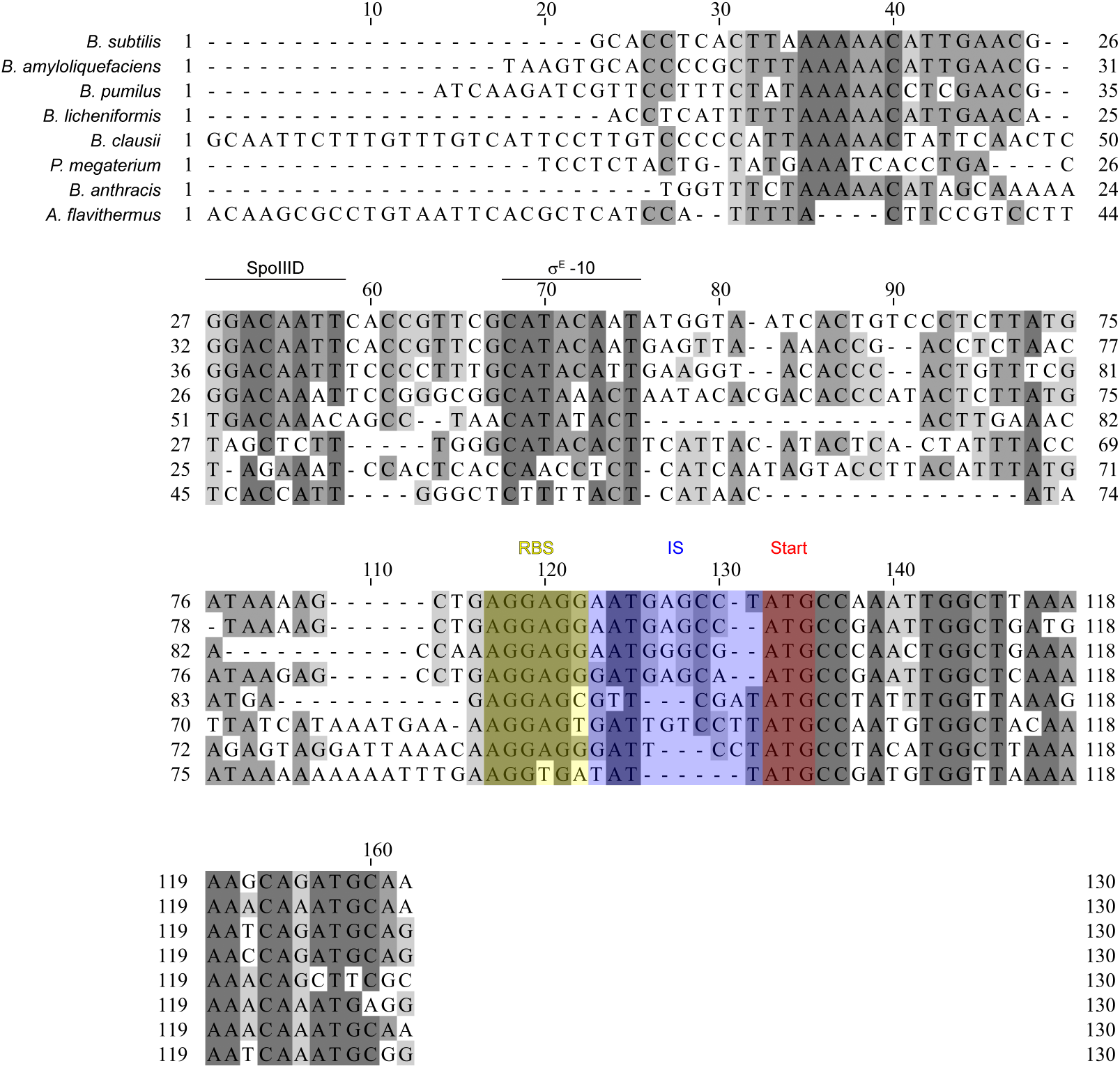
Sequence conservation of the *cmpA* RBS, IS, and first several codons of the open reading frame across related species. Sequence conservation of the upstream region and first 10 codons of *cmpA* across various species: *Bacillus subtilis* (NC_000964.3), *Bacillus amyloliquefaciens* (NZ_CP072120.1), *Bacillus pumilus* (NC_009848.4), *Bacillus licheniformis* (NZ_CP140161.1), *Bacillus clausii* (NZ_CP140150.1), *Priestia megaterium* (NZ_CP035094.1), *Bacillus anthracis* (NC_007530.2), and *Anoxybacillus flavithermus* (NC_011567.1). The RBS is highlighted in yellow, the IS is highlighted in blue, and the start codon is highlighted in red.

**Figure S6.**
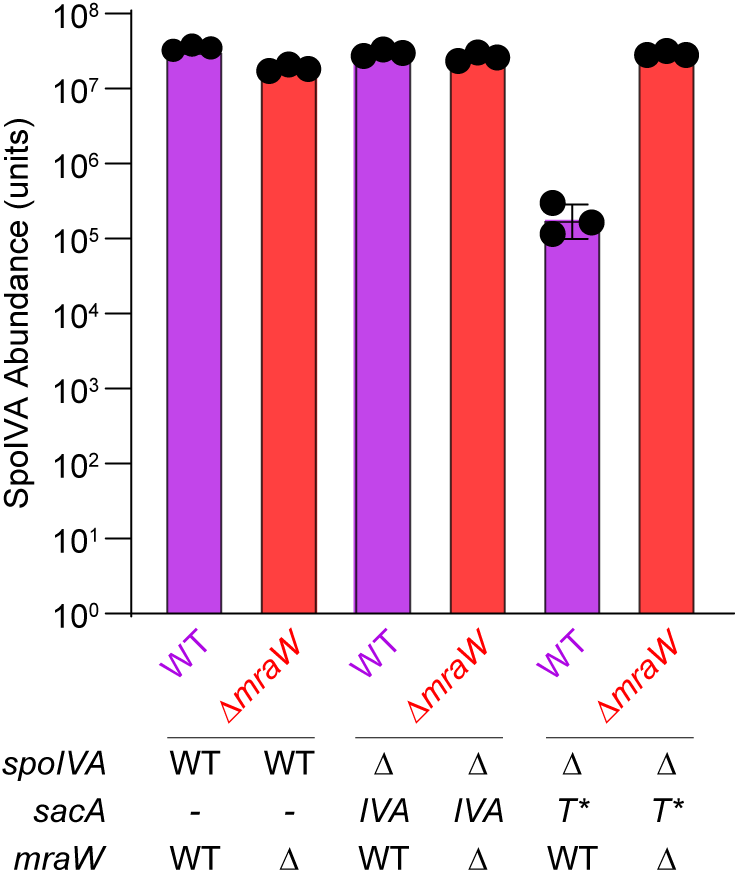
SpoIVA^T*^ levels are restored to wild type levels in Δ*mraW*. SpoIVA abundance measured by mass-spectrometry in strains harboring the indicated allele of *spoIVA* in the presence and absence of *mraW*. *sacA* is a chromosomal locus from which the indicated allele of *spoIVA* is expressed. Strains: PY79, ZP1, ZP429, ZP435, ZP379, and ZP381. Bars represent mean (n = 3 biological replicates); errors: S.D.

**Figure S7.**
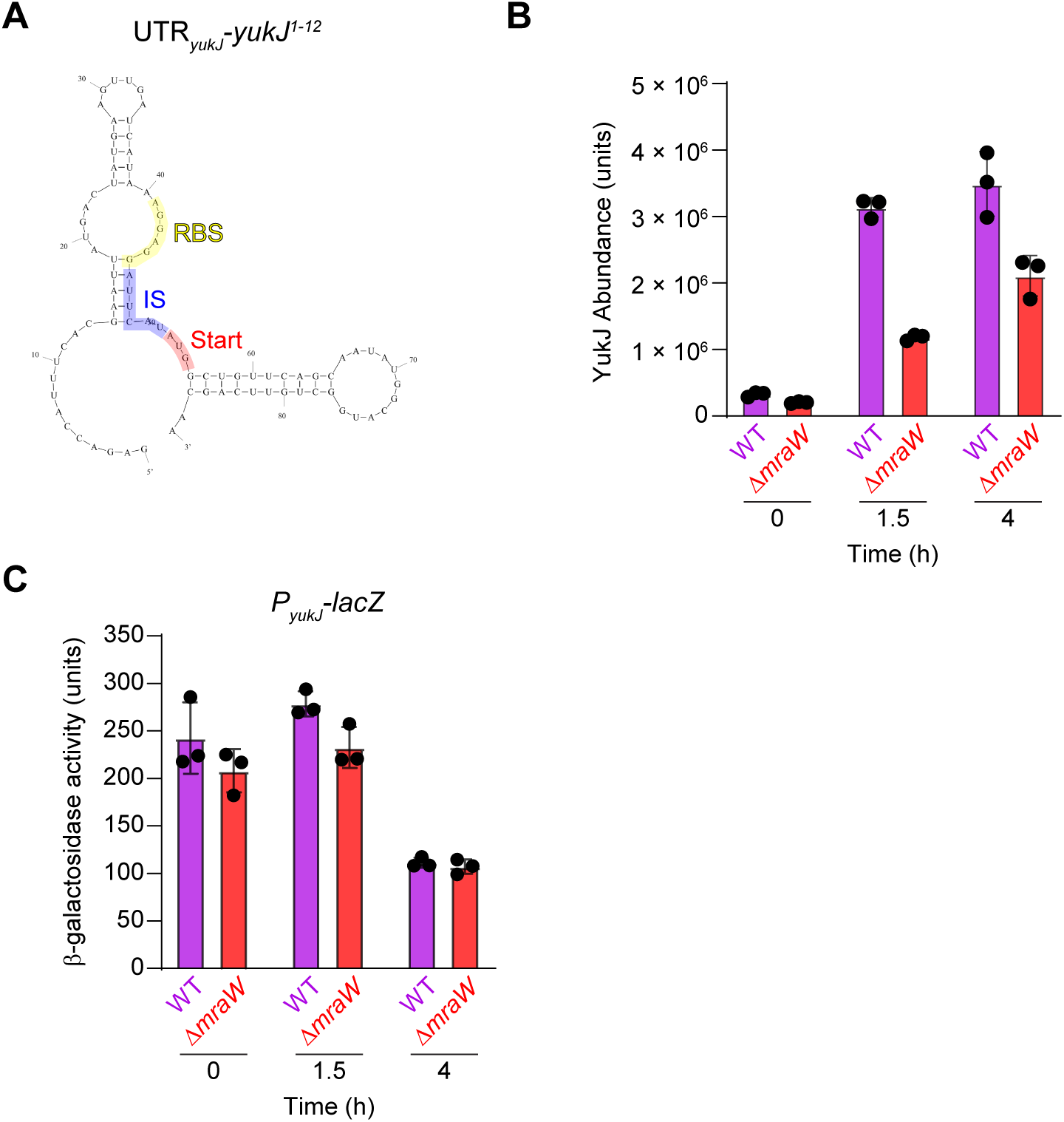
YukJ translation is dependent on MraW-modified ribosomes. (A) Mfold predicted structure of the 5’UTR and first 12 codons of *yukJ*. (B) YukJ abundance measured by mass spectrometry in the presence (purple) or absence (red) of *mraW* at the indicated time points. Bars represent mean (n = 3 biological replicates); errors: S.D. Strains: PY79 and ZP1. (C) β-galactosidase activity arising from a transcriptional fusion of the promoter of *yukJ* to *lacZ* in the presence (purple) or absence (red) of *mraW* at the indicated time points. Bars represent mean (n = 3 biological replicates); errors: S.D. Strains: ZP546 and ZP550.

**Table S1.** *B. subtilis* strains used in this study.

| Strain | Genotype | Reference |
| --- | --- | --- |
| PY79 | Prototrophic derivative of *B. subtilis* 168 | 1 |
| KP73 | $\Delta spoIVA::kan$ | 2 |
| IT895 | $\Delta spoIVA::kan \Delta cmpA::erm$ | 3 |
| JPC282 | $\Delta spoIVA::erm \Delta spoVID::kan$ | This study |
| ZP44 | $\Delta spoIVA::kan \Delta mraW::erm$ | This study |
| KR394 | $\Delta spoIVA::kan thrC::spoIVA spec$ | 4 |
| ZP47 | $\Delta spoIVA::kan thrC::spoIVA^{L59P} spec$ | This study |
| ZP50 | $\Delta spoIVA::kan \Delta cmpA::erm thrC::spoIVA^{L59P} spec$ | This study |
| ZP53 | $\Delta spoIVA::erm \Delta spoVID::kan thrC::spoIVA^{L59P} spec$ | This study |
| ZP56 | $\Delta spoIVA::kan \Delta mraW::erm thrC::spoIVA^{L59P} spec$ | This study |
| KR160 | $thrC::gfp-spoIVA spec$ | 4 |
| ZP35 | $\Delta spoIVA::kan amyE::spoIVA^{L59P} cat thrC::gfp-spoIVA^{L59P} spec$ | This study |
| ZP77 | $\Delta spoIVA::kan thrC::spoIVA spec \Delta mraW::erm$ | This study |
| KR367 | $\Delta spoIVA::kan thrC::spoIVA^{K30A} spec$ | 3 |
| ZP62 | $\Delta spoIVA::kan \Delta mraW::erm thrC::spoIVA^{K30A} spec$ | This study |
| JPC221 | $\Delta spoIVA::kan thrC::spoIVA^{T70A, T71A} spec$ | 5 |
| ZP65 | $\Delta spoIVA::kan \Delta mraW::erm thrC::spoIVA^{T70A, T71A} spec$ | This study |
| JB103 | $\Delta spoIVA::kan thrC::spoIVA^{D97A} spec$ | 6 |
| ZP59 | $\Delta spoIVA::kan \Delta mraW::erm thrC::spoIVA^{D97A} spec$ | This study |
| ZP134 | $\Delta spoIVA::kan thrC::spoIVA spec \Delta mraW::erm amyE::mraW cat$ | This study |
| ZP231 | $\Delta spoIVA::kan thrC::spoIVA spec \Delta mraW::erm amyE::mraW^{D52A} cat$ | This study |
| ZP233 | $\Delta spoIVA::kan thrC::spoIVA spec \Delta mraW::erm amyE::mraW^{D100A} cat$ | This study |
| ZP137 | $\Delta spoIVA::kan \Delta mraW::erm thrC::spoIVA^{L59P} spec amyE::mraW cat$ | This study |
| ZP239 | $\Delta spoIVA::kan thrC::spoIVA^{L59P} spec \Delta mraW::erm amyE::mraW^{D52A} cat$ | This study |
| ZP241 | $\Delta spoIVA::kan thrC::spoIVA^{L59P} spec \Delta mraW::erm amyE::mraW^{D100A} cat$ | This study |
| ZP221 | $\Delta spoIVA::kan thrC::spoIVA spec \Delta yabC::erm$ | This study |
| ZP287 | $\Delta spoIVA::kan \Delta mraW::erm thrC::spoIVA spec \Delta yabC::erm::cat$ | This study |
| ZP215 | $\Delta spoIVA::kan thrC::spoIVA^{L59P} spec \Delta yabC::erm$ | This study |
| ZP283 | $\Delta spoIVA::kan \Delta mraW::erm thrC::spoIVA^{L59P} spec \Delta yabC::erm::cat$ | This study |
| ZP249 | $\Delta spoIVA::kan thrC::spoIVA^{T70A, T71A} spec \Delta yabC::erm$ | This study |
| ZP285 | $\Delta spoIVA::kan \Delta mraW::erm thrC::spoIVA^{T70A, T71A} spec \Delta yabC::erm::cat$ | This study |
| ZP1 | $\Delta mraW::erm$ | This study |
| ZP89 | $\Delta spoIVA::kan \Delta mraW::erm thrC::spoIVA spec amyE::P_{spoIVA-} mraW cat$ | This study |
| ZP83 | $\Delta spoIVA::kan \Delta mraW::erm thrC::spoIVA^{L59P} spec amyE::P_{spoIVA-mraW} cat$ | This study |
| ZP107 | $\Delta spoIVA::kan \Delta mraW::erm thrC::spoIVA spec amyE::P_{cotR-mraW} cat$ | This study |
| ZP104 | $\Delta spoIVA::kan \Delta mraW::erm thrC::spoIVA^{L59P} spec amyE::P_{cotR-mraW} cat$ | This study |
| ZP385 | $\Delta spoIVA::kan sacA::spoIVA^{T70A, T71A} cat::spec amyE::P_{cmpA-cmpA-lacZ} cat$ | This study |
| ZP389 | $\Delta spoIVA::kan 1::spoIVA^{T70A, T71A} cat::spec \Delta mraW::erm amyE::P_{cmpA-cmpA-lacZ} cat$ | This study |
| ZP383 | $\Delta spoIVA::kan sacA::spoIVA^{T70A, T71A} cat::spec amyE::P_{cmpA-lacZ} cat$ | This study |
| ZP387 | $\Delta spoIVA::kan sacA::spoIVA^{T70A, T71A} cat::spec \Delta mraW::erm amyE::P_{cmpA-lacZ} cat$ | This study |
| SE230 | $amyE::P_{cmpA-cmpA-lacZ} cat$ | 7 |
| ZP313 | $amyE::P_{cmpA-cmpA-lacZ} cat \Delta mraW::erm$ | This study |
| SE222 | $amyE::P_{cmpA-lacZ} cat$ | 7 |
| ZP311 | $amyE::P_{cmpA-lacZ} cat \Delta mraW::erm$ | This study |
| IT686 | $sacA::P_{sigA-cmpA} cat$ | 3 |
| ZP116 | $\Delta mraW::erm sacA::P_{sigA-cmpA} cat$ | This study |
| ZP315 | $\Delta spoIVA::kan thrC::spoIVA spec sacA::P_{sigA-cmpA} cat$ | This study |
| ZP321 | $\Delta spoIVA::kan thrC::spoIVA spec amyE::P_{sigA-cmpA} cat \Delta mraW::erm$ | This study |
| ZP317 | $\Delta spoIVA::kan thrC::spoIVA^{T70A, T71A} spec amyE::P_{sigA-cmpA} cat$ | This study |
| ZP319 | $\Delta spoIVA::kan thrC::spoIVA^{T70A, T71A} spec amyE::P_{sigA-cmpA} cat \Delta mraW::erm$ | This study |
| ZP441 | $amyE::P_{cmpA-cmpA^{1-30}-lacZ} cat$ | This study |
| ZP446 | $amyE::P_{cmpA-cmpA^{1-30}-lacZ} cat \Delta mraW::erm$ | This study |
| ZP439 | $amyE::P_{cmpA-cmpA^{1-20}-lacZ} cat$ | This study |
| ZP444 | $amyE::P_{cmpA-cmpA^{1-20}-lacZ} cat \Delta mraW::erm$ | This study |
| ZP437 | $amyE::P_{cmpA-cmpA^{1-10}-lacZ} cat$ | This study |
| ZP442 | $amyE::P_{cmpA-cmpA^{1-10}-lacZ} cat \Delta mraW::erm$ | This study |
| ZP463 | $amyE::P_{cmpA-cmpA^{1-20} (-1 frameshift)-lacZ} cat$ | This study |
| ZP469 | $amyE::P_{cmpA-cmpA^{1-20} (-1 frameshift)-lacZ} cat \Delta mraW::erm$ | This study |
| ZP560 | $sacA::P_{sigA-IS^*-cmpA} cat$ | This study |
| ZP562 | $sacA::P_{sigA-IS^*-cmpA} cat \Delta mraW::erm$ | This study |
| ZP451 | $amyE::P_{cmpA-cmpA^{1-4}-lacZ} cat$ | This study |
| ZP457 | $amyE::P_{cmpA-cmpA^{1-4}-lacZ} cat \Delta mraW::erm$ | This study |
| ZP473 | $amyE::P_{cmpA-cmpA^{1-6}-lacZ} cat$ | This study |
| ZP481 | $amyE::P_{cmpA-cmpA^{1-6}-lacZ} cat \Delta mraW::erm$ | This study |
| ZP503 | $amyE::P_{cmpA-cmpA^{stem-swap}-lacZ} cat$ | This study |
| ZP497 | $amyE::P_{cmpA-cmpA^{stem-swap}-lacZ} cat \Delta mraW::erm$ | This study |
| ZP532 | $amyE::P_{cmpA-cmpA^{1-6} (codons 4*/5*)-lacZ} cat$ | This study |
| ZP534 | $amyE::P_{cmpA-cmpA^{1-6} (codons 4*/5*)-lacZ} cat \Delta mraW::erm$ | This study |
| ZP568 | *amyE::P<sub>cmpA</sub>-cmpA<sup>1-6</sup> (stem strong)-lacZ cat* | This study |
| ZP570 | *amyE::P<sub>cmpA</sub>-cmpA<sup>1-6</sup> (stem strong)-lacZ cat ΔmraW::erm* | This study |
| ZP507 | *amyE::P<sub>isp</sub>-lacZ cat* | This study |
| ZP513 | *amyE::P<sub>isp</sub>-lacZ cat ΔmraW::erm* | This study |
| ZP509 | *amyE::P<sub>isp</sub> (stem-swap)-lacZ cat* | This study |
| ZP515 | *amyE::P<sub>isp</sub> (stem-swap)-lacZ cat ΔmraW::erm* | This study |
| ZP577 | *amyE::P<sub>isp</sub> (IS\*)-lacZ cat* | This study |
| ZP583 | *amyE::P<sub>isp</sub> (IS\*)-lacZ cat ΔmraW::erm* | This study |
| ZP572 | *amyE::P<sub>spoIIQ</sub>-T7 pol spec sacA::P<sub>T7</sub>-NEB5'UTR-gfp cat* | This study |
| ZP574 | *amyE::P<sub>spoIIQ</sub>-T7 pol spec sacA::P<sub>T7</sub>-NEB5'UTR-gfp cat ΔmraW::erm* | This study |
| ZP405 | *ΔspoIVA::kan sacA::spoIVA<sup>T70A, T71A</sup> cat::spec amyE::P<sub>spoVID</sub>-spoVID-lacZ cat* | This study |
| ZP410 | *ΔspoIVA::kan sacA::spoIVA<sup>T70A, T71A</sup> cat::spec amyE::P<sub>spoVID</sub>-spoVID-lacZ cat ΔmraW::erm* | This study |
| ZP339 | *amyE::P<sub>spoVID</sub>-spoVID-lacZ cat* | This study |
| ZP345 | *amyE::P<sub>spoVID</sub>-spoVID-lacZ cat ΔmraW::erm* | This study |
| ZP363 | *amyE::P<sub>sigA</sub>-lacZ cat* | This study |
| ZP361 | *amyE::P<sub>sigA</sub>-lacZ cat ΔmraW::erm* | This study |
| ZP523 | *amyE::P<sub>sigA</sub>-cmpA-gfp cat* | This study |
| ZP525 | *amyE::P<sub>sigA</sub>-cmpA-gfp cat ΔmraW::erm* | This study |
| ZP429 | *ΔspoIVA::kan sacA::spoIVA cat::spec* | This study |
| ZP435 | *ΔspoIVA::kan sacA::spoIVA cat::spec ΔmraW::erm* | This study |
| ZP379 | *ΔspoIVA::kan sacA::spoIVA<sup>T70A, T71A</sup> cat::spec* | This study |
| ZP381 | *ΔspoIVA::kan sacA::spoIVA<sup>T70A, T71A</sup> cat::spec ΔmraW::erm* | This study |
| ZP546 | *amyE::P<sub>yukJ</sub>-lacZ cat* | This study |
| ZP550 | *amyE::P<sub>yukJ</sub>-lacZ cat ΔmraW::erm* | This study |
| ZP511 | *amyE::P<sub>isp</sub> (stem strong)-lacZ cat* | This study |
| ZP517 | *amyE::P<sub>isp</sub> (stem strong)-lacZ cat ΔmraW::erm* | This study |

